# Coordinated activity and plasticity of infralimbic cortex GABAergic interneurons are critical for fear extinction encoding

**DOI:** 10.64898/2026.02.07.704584

**Authors:** Rodrigo Campos-Cardoso, Hunter T. Franks, Noelle R. Potter, Zephyr R. Desa, Brianna L. Fitzgerald, Rebekah L. Fowler, Faith A. Brown, Victoria A. Landar, Meris Privette, Kirstie A. Cummings

**Author notes:** These authors contributed equally (Hunter T. Franks, Noelle R. Potter). Correspondence: Kirstie A. Cummings.

## Abstract

The ability for an organism to encode fear memories is necessary for survival. Once a threat is no longer present, organisms must suppress, or extinguish, this fear memory in favor of other adaptive behaviors. In rodents, the infralimbic cortex (IL) is a locus critical for the extinction of cued fear memory. While this role has been known for decades, the circuit mechanisms underlying its recruitment are largely unknown. By using a combination of immunohistochemistry, neural tagging, *in vivo* calcium imaging and optogenetics, and optogenetics-assisted brain slice electrophysiology, we revealed that the dynamic activity and plasticity of IL inhibitory interneurons is critical for encoding fear extinction. Specifically, after fear conditioning, IL parvalbumin interneurons exhibit increased activity and plasticity, driving enhanced freezing. After fear extinction, however, IL somatostatin interneurons exhibit extinction cue-associated activity and plasticity, and their activity facilitates extinction memory encoding through inhibition of parvalbumin interneuron activity and disinhibition of IL principal neurons. Further, glutamatergic projections from the basolateral amygdala undergo experience- and cell type-specific plasticity that is required to drive the dynamic recruitment of IL parvalbumin and somatostatin interneurons after fear conditioning and extinction, respectively. Overall, these results reveal the mechanisms of cued fear extinction encoding and highlight critical roles for local IL microcircuit computations in these roles.

## Introduction

Fear memory encoding is critical for the ability of an organism to identify and appropriately respond to threats, ultimately promoting survival^1,2^. Equally as important is for organisms to learn to suppress fear memories in the absence of threats in order to resume other adaptive behaviors. Dysfunction of fear suppression (i.e., fear extinction) is central to several neuropsychiatric disorders like post-traumatic stress disorder^3–5^. Compared to the circuits of fear memory encoding, however, the circuits underlying the extinction of fear memory are less well-understood.

The infralimbic (IL) subregion of the rodent medial prefrontal cortex (mPFC) is critical for the extinction of auditory fear memory^6–9^. Foundational studies that manipulate IL activity during extinction training via pharmacological, lesioning, electrical stimulation, and optogenetic methods result in alterations in extinction memory retrieval the next day, underscoring the critical role for IL in extinction encoding^10–15^. IL activity during extinction retrieval is also important for extinction memory expression. *In vivo* electrophysiological recordings and immunohistochemical experiments revealed that increases in IL neural activity are also correlated with retrieval of extinction memory^16–19^. Once recruited, IL principal projection neurons (PNs) engage basolateral (BLA) and basomedial amygdala to promote extinction behaviors^20–25^. However, the local and long-range circuit mechanisms that drive the recruitment of IL PNs and subsequent extinction encoding are largely unstudied.

While generally understudied, some studies have revealed important roles for mPFC interneurons in fear memory processes. In the prelimbic subregion of mPFC, the activity and plasticity of local inhibitory interneurons is required for fear memory encoding^26,27^. While much less is known about whether and how these cells may be dictating fear extinction encoding in IL, some studies hint towards potential roles. Specifically, stress-induced deficits in fear extinction are driven at least in part by IL interneurons^28^. Additionally, IL interneurons in adolescent rodents have been implicated in the regulation of fear extinction^29^. Despite these clues for potential roles for interneurons in fear extinction processing, the mechanisms of such roles are unknown.

Here, we sought to reveal the mechanisms by which local IL GABAergic microcircuits shape cued fear memory extinction. We used a combination of neural tagging, immunohistochemistry, calcium imaging, *in vivo* optogenetics, and *ex vivo* brain slice electrophysiology to reveal that the dynamic activity and plasticity of IL GABAergic circuits is critical for the extinction of auditory fear memory. These results underscore the important yet largely underappreciated roles that GABAergic circuits play in shaping multiple facets of emotional memory.

## Results

While IL PN activity is critical for fear extinction memory processes, how local GABAergic interneurons are recruited to shape activity within this region is less well-understood. We therefore performed cFos immunohistochemistry in PV-Cre/ Ai9 and SST-Flp/ Ai65F transgenic mouse lines that specifically fluorescently label the two most abundant cortical interneuron populations, parvalbumin (PV-INs) and somatostatin (SST-INs) -expressing interneurons^30,31^. Mice underwent either auditory tone conditioned stimulus (CS) exposure followed by CS tone re-exposure in a distinct context 24 hours later (tones group; **Fig. 1a**); auditory fear conditioning consisting of 6 pairings of CS and co-terminating foot shock followed by CS-evoked fear memory retrieval in a distinct context 24 hours later (conditioning group; **Fig. 1b**); or paired auditory fear conditioning and two consecutive days of auditory fear extinction training consisting of 20 CS presentations in a distinct context followed by CS-evoked extinction memory retrieval 24 hours later (extinction group; **Fig. 1c**) (**Supplemental Fig. 1**). At 90 minutes after tone re-exposure or fear or extinction memory retrieval, brains were isolated, sectioned, and immunostained for cFos (**Fig. 1c and 1d**). cFos puncta and their overlap with fluorescently labeled IL SST-INs and PV-INs were then quantified. Increased cFos density was observed in IL in the extinction group compared to tones and conditioned groups, consistent with our previous results and other studies (**Fig 1g,k**)^19,32^. While there were no differences in the density of IL SST-INs across groups (**Fig. 1f**), there was an increase and decrease in the density of cFos+ IL SST-INs in the extinction and conditioning groups, respectively (**Fig. 1h**). Normalization of the number of cFos+ SST-INs to the total number of IL SST-INs in each group revealed the same effects (**Fig. 1i**). The same quantification was performed for IL PV-INs. While density of IL PV-INs was not different across behavior groups (**Fig. 1j**), we observed an increase in the density (**Fig. 1l**) and proportion (**Fig. 1m**) of cFos+ IL PV-INs in the conditioning group compared to tones or extinction groups. These results indicate that IL SST-INs and PV-INs are dynamically recruited across relative low and high fear states, suggesting they may play distinct roles in these processes.

**Figure 1.**
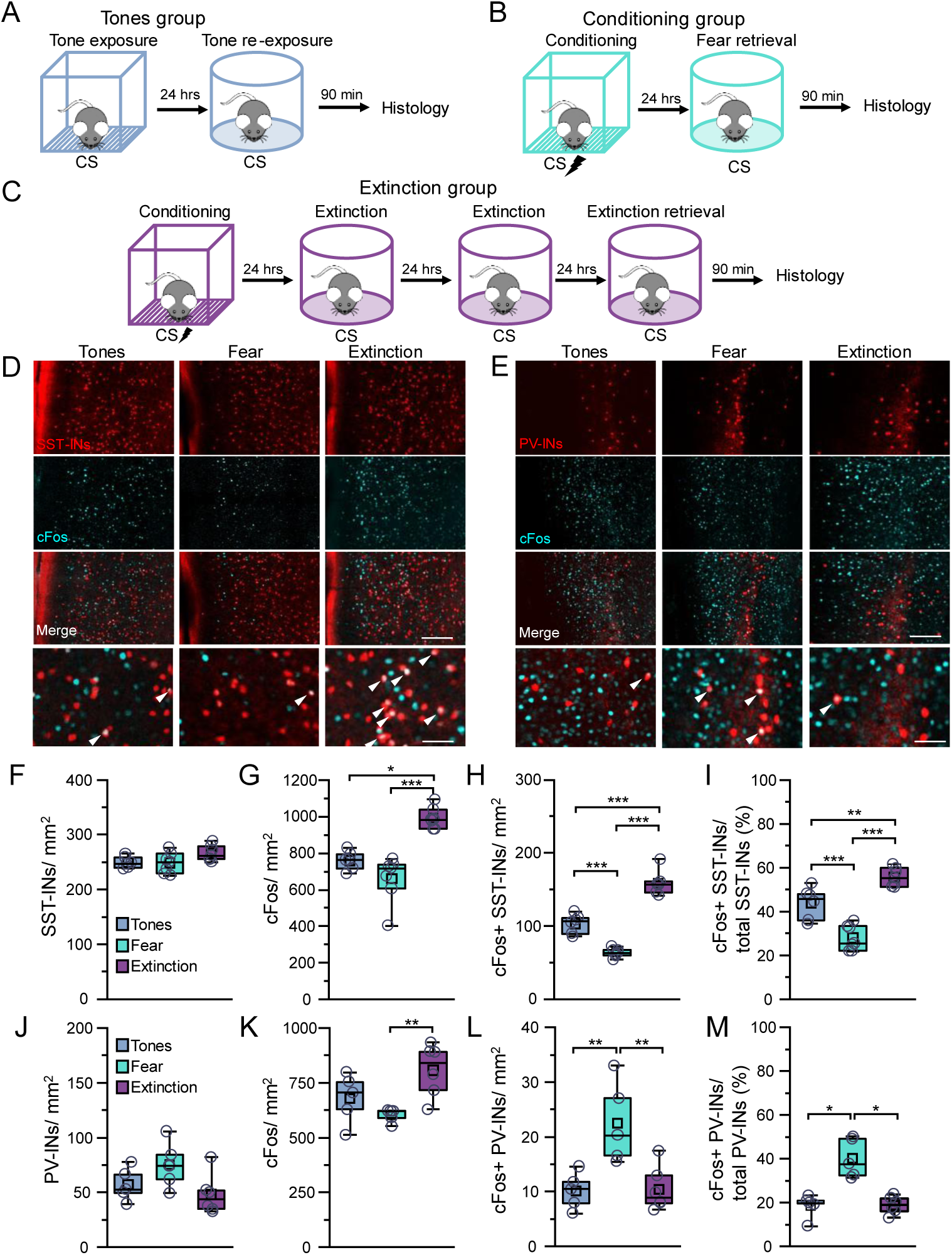
Fear and extinction memory retrieval drive differential recruitment of infralimbic cortex SST-INs and PV-INs. Mice were subjected to (A) six auditory tones (2 kHz, 80 db, 20 s) followed by a tone re-exposure test 24 hours later (4 tones, 2 kHz, 80 db, 20 s); (B) six pairings of auditory tones (2 kHz, 80 db, 20 s) and co-terminating foot shocks (0.7 mA, scrambled, 2 s) followed by a tone-evoked fear memory retrieval test (4 tones, 2 kHz, 80 db, 20 s) 24 hours later; or (C) six pairings of auditory tones (2 kHz, 80 db, 20 s) and co-terminating foot shocks (0.7 mA, scrambled, 2 s) followed by two consecutive days of auditory fear memory extinction training (20 tones, 2 kHz, 80 db, 20 s) and a tone-evoked fear extinction memory retrieval test (4 tones, 2 kHz, 80 db, 20 s) 24 hours after extinction training. At 90 minutes after tone re-exposure, fear memory retrieval, or extinction memory retrieval tests, mice were subjected to transcardial perfusion for cFos immunohistochemistry. (D-E) Representative histological sections of infralimbic cortex from (D) SST-Flp/ Ai65F and (E) PV-Cre/ Ai9 mice from the tones group, conditioning group, or extinction group. Scale = 100 µm (or 25 µm for lower panels). Quantification of (F) the density of SST-INs (F_2,17_ = 3.10, p = 0.07, one-way ANOVA), (G) cFos (Χ^2^ = 14.27 [2], p = 7.95×10^-4^, Kruskal-Wallis ANOVA), (H) cFos^+^ SST-INs (F_2,17_ = 104.2, p = 0.0001, one-way ANOVA), and (I) proportion of all SST-INs that are cFos^+^ (F_2,17_ = 39.80, p = 0.0001, one-way ANOVA) in infralimbic cortex of SST-Flp/ Ai65F mice subjected to tones-only (n = 6 mice, 18 slices), conditioning (n = 7 mice, 21 slices), and extinction (n = 7 mice, 21 slices). Quantification of (J) the density of PV-INs (F_2,13_ = 2.96, p = 0.09, one-way ANOVA), (K) cFos (Χ^2^ = 5.16 [2], p = 0.04, Kruskal-Wallis ANOVA), (L) cFos^+^ PV-INs (F_2,13_ = 9.56, p = 0.003, one-way ANOVA), and (M) proportion of all PV-INs that are cFos^+^ (Χ^2^ = 9.71 [2], p = 0.008, Kruskal-Wallis ANOVA) in infralimbic cortex of PV-Cre/ Ai9 mice subjected to tones-only (n = 5 mice, 15 slices), conditioning (n = 5 mice, 15 slices), and extinction (n = 6 mice, 18 slices). *, p < 0.05, **, p < 0.01, and ***, p < 0.001, Tukey’s or Dunn’s post-hoc test. Boxplots represent the median (center line), mean (square), quartiles, and 10%–90% range (whiskers/error bars). Open circles represent data points for individual mice.

We next tested how IL SST-INs and PV-INs are recruited to fear- or extinction-tagged neural ensembles. We used an AAV vector encoding ERCreER under the control of an engineered activity-dependent promoter fused to *arc min* (ESARE-ERCreER)^33^ along with a Cre-dependent eYFP vector and a pan-neuronal mCherry vector to allow us to estimate the extent of viral transduction in IL (**Fig. 2a**). Upon intraperitoneal injection of 4-hydroxytamoxifen (4-OHT), active neurons can be permanently tagged with eYFP. All mice were subjected to paired fear conditioning after which they were split randomly into one of three groups and subjected to additional tests 24 hours after. The first group was subjected to a CS-evoked fear memory retrieval test (fear retrieval group); the second group was subjected to one day of fear extinction (early extinction group); and the third group was subjected to 3 consecutive days of fear extinction, each separated by 24 hours (late extinction group) (**Fig. 2b, Supplemental Fig. 2**). Immediately after the last training session in each group, mice received intraperitoneal injections of 4-OHT. Brain sections were procured across groups and stained for somatostatin (**Fig. 2c**) and parvalbumin (**Fig. 2i**) protein. First, we quantified the number of IL neurons recruited across groups (eYFP^+^ cells) and observed an increased in late extinction compared to either fear retrieval or early extinction (**Fig 2d, 2j**). This observation is in line with the idea that IL is broadly recruited in response to fear extinction, consistent with our previous data (**Fig. 1**) and the literature^19,32^. While the numbers of IL SST-INs were not different across groups (**Fig. 2e**), they were more highly recruited to the IL neural ensemble in late extinction compared to early extinction or fear retrieval (**Fig. 2f**). In turn, the proportion SST+ of neurons in the tagged ensembles increased across extinction (**Fig 2g**), as did the proportion of total IL SST-INs that were recruited to the IL ensemble (**Fig. 2h**). IL PV-IN counts were not different across groups (**Fig. 2k**). In contrast to SST-INs, we observed the highest recruitment of PV-INs following fear memory retrieval, which decreased across extinction (**Fig. 2l**). In line with these results, the proportion of PV-INs comprising the IL neural ensemble decreased across extinction (**Fig. 2m**), as did the proportion of total IL PV-INs that were recruited to the IL ensemble across extinction (**Fig. 2n**). As the ESARE vector relies on an *arc*-based promoter, we also independently confirmed these patterns of activity by using two different readouts of neural activity (cFos- and Arc-based approaches). Overall, these results indicate that IL SST-INs are recruited during extinction, whereas IL PV-INs are recruited in response to fear and are disengaged in response to extinction training.

**Figure 2.**
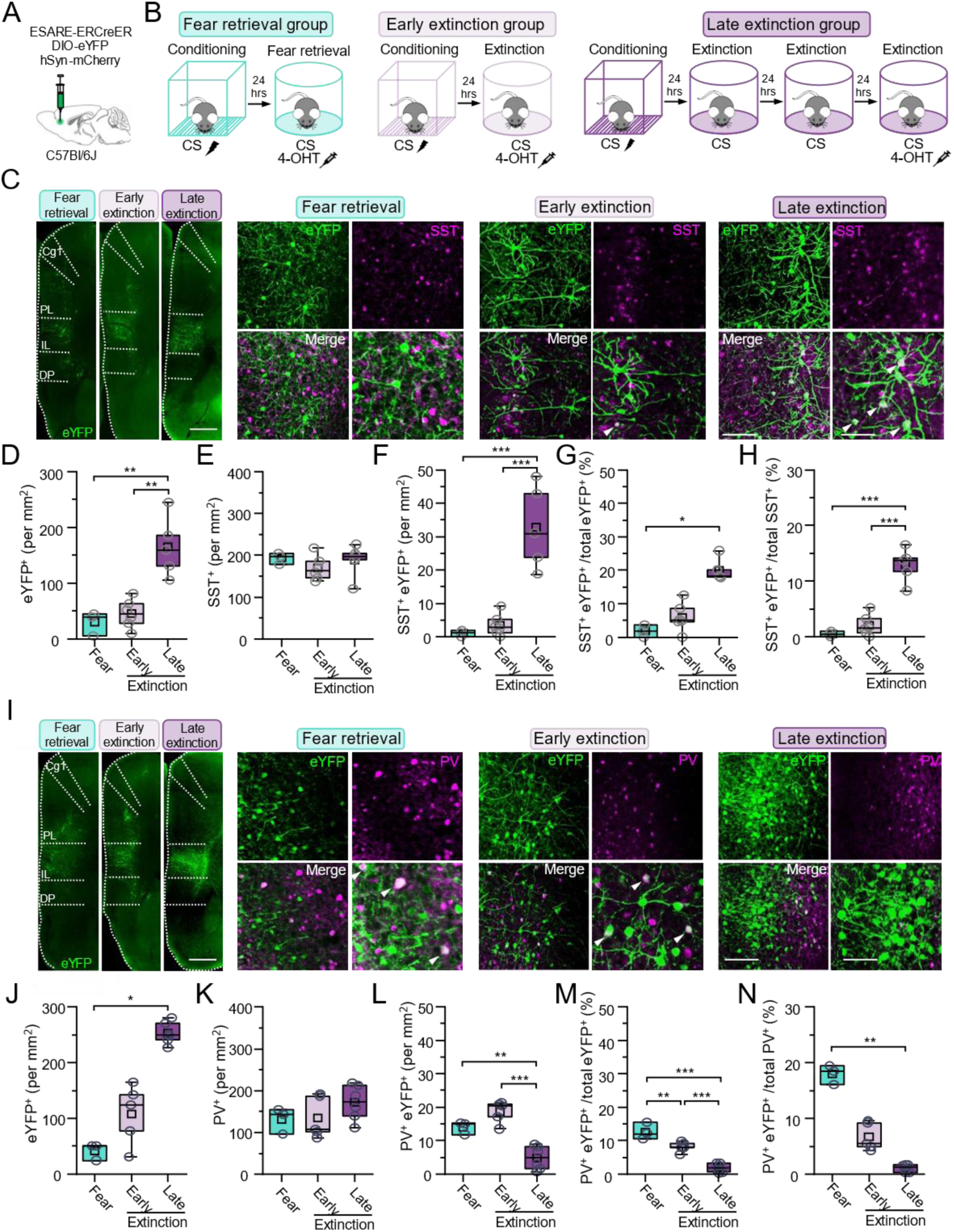
Infralimbic cortex PV-INs and SST-INs are differentially recruited to fear- and extinction-related neural ensembles. (A) Wild-type mice received infusions of a cocktail of ESARE-ERCreER, DIO-eYFP, and hSyn-mCherry in the infralimbic cortex and returned to the home cage for 3 weeks. (B) All mice were first subjected to six pairings of auditory tones (2 kHz, 80 db, 20 s) and co-terminating foot shocks (0.7 mA, scrambled, 2 s). The fear retrieval group was then subjected to a tone-evoked fear memory retrieval test (4 tones, 2 kHz, 80 db, 20 s) 24 hours later (Fear retrieval group). The early and late extinction groups were subjected to one or three consecutive days of auditory fear extinction training (20 tones per session, 2 kHz, 80 db, 20 s), respectively. Immediately the last behavioral session for each group, mice received a 10 mg/kg intraperitoneal injection of 4-hydroxytamoxifen (4-OHT). Two weeks later, mice were subjected to transcardial perfusion for immunohistochemical analysis. (C) Representative infralimbic cortex sections containing tagged neurons and immunohistochemically stained SST-Ins from mice in the fear retrieval, early extinction, or late extinction groups. Scale = 500 µm (100 µm in panels and 50 µm in zoom-in). Quantification of (D) the density of eYFP^+^ neurons (F_2,11_ = 17.63, p = 3.71×10^-4^, one-way ANOVA), (E) SST^+^ neurons (F_2,11_ = 0.76, p = 0.49, one-way ANOVA), (F) eYFP^+^/SST^+^ double positive neurons (Χ^2^ = 9.89 [2], p = 0.007, Kruskal-Wallis ANOVA), (G) proportion of total eYFP^+^ neurons that are SST^+^ (Χ^2^ = 10.23 [2], p = 0.006, Kruskal-Wallis ANOVA), and (H) proportion of total SST^+^ neurons that are eYFP^+^ (F_2,11_ = 40.63, p = 0.0001, one-way ANOVA) in infralimbic cortex of mice from fear retrieval (n = 3 mice, 9 slices), early extinction (n = 6 mice, 18 slices), and late extinction (n = 5 mice, 15 slices) groups. (I) Representative infralimbic cortex sections containing tagged neurons and immunohistochemically stained PV-INs from mice in the fear retrieval, early extinction, or late extinction groups. Scale = 500 µm (100 µm in panels and 50 µm in zoom-in). Quantification of (J) the density of eYFP^+^ neurons (Χ^2^ = 8.70 [2], p = 0.013, Kruskal-Wallis ANOVA), (K) PV^+^ neurons (F_2,11_ = 2.71, p = 0.11, one-way ANOVA), (L) eYFP^+^/PV^+^ double positive neurons (F_2,11_ = 26.88, p = 0.0001, one-way ANOVA), (M) proportion of total eYFP^+^ neurons that are PV^+^ (F_2,11_ = 45.6, p = 0.0001, one-way ANOVA), (N) and proportion of total PV^+^ neurons that are eYFP^+^ (Χ^2^ = 11.3 [2], p = 0.003, Kruskal-Wallis ANOVA) in infralimbic cortex of mice belonging to fear retrieval (n = 3 mice, 9 slices), early extinction (n = 6 mice, 18 slices), and late extinction (n = 5 mice, 15 slices) groups. *, p < 0.05, **, p < 0.01, and ***, p < 0.001, Tukey’s or Dunn’s post-hoc test. Boxplots represent the median (center line), mean (square), quartiles, and 10%–90% range (whiskers/error bars). Open circles represent data points for individual mice.

While the results from above suggest differential recruitment of IL SST-INs and PV-INs across distinct fear and extinction states, all readouts are post-training and represent a single timepoint. We therefore performed *in vivo* calcium imaging to observe when IL SST-INs and PV-INs are recruited and how their activity evolves in real time across training in awake behaving mice. SST-Cre or PV-Cre mice received unilateral infusions of a Cre-dependent GCaMP8f into IL along with imaging fiber implantation (**Fig 3a, Supplementary Fig. 3**). Calcium signals were recorded continuously during a pre-conditioning tone exposure test, fear conditioning, fear memory retrieval, extinction days 1 and 2, and extinction memory retrieval and analyzed and compared to behavior across sessions in the same mice (**Supplementary Fig. 4**). Compared to fear memory retrieval, IL SST-IN activity increased across extinction learning and retrieval (**Fig. 3b-d**). In contrast, IL PV-IN activity decreased across extinction learning and retrieval compared to fear memory retrieval (**Fig. 3e-g**). We performed an additional analysis from a random subset of the same animals during which mice were freezing during baseline or the intertrial intervals (epochs between CS presentations) in all behavior tests. These analyses revealed that the activity of IL SST-INs and PV-INs did not simply signal ambulation or freezing *per se* (**Supplemental Fig. 5**). Rather, IL interneurons specifically signal fear- or extinction-related information during CS presentation. These results are in line with the idea that IL SST-INs and PV-INs are dynamically and orthogonally engaged across low and high fear states.

**Figure 3.**
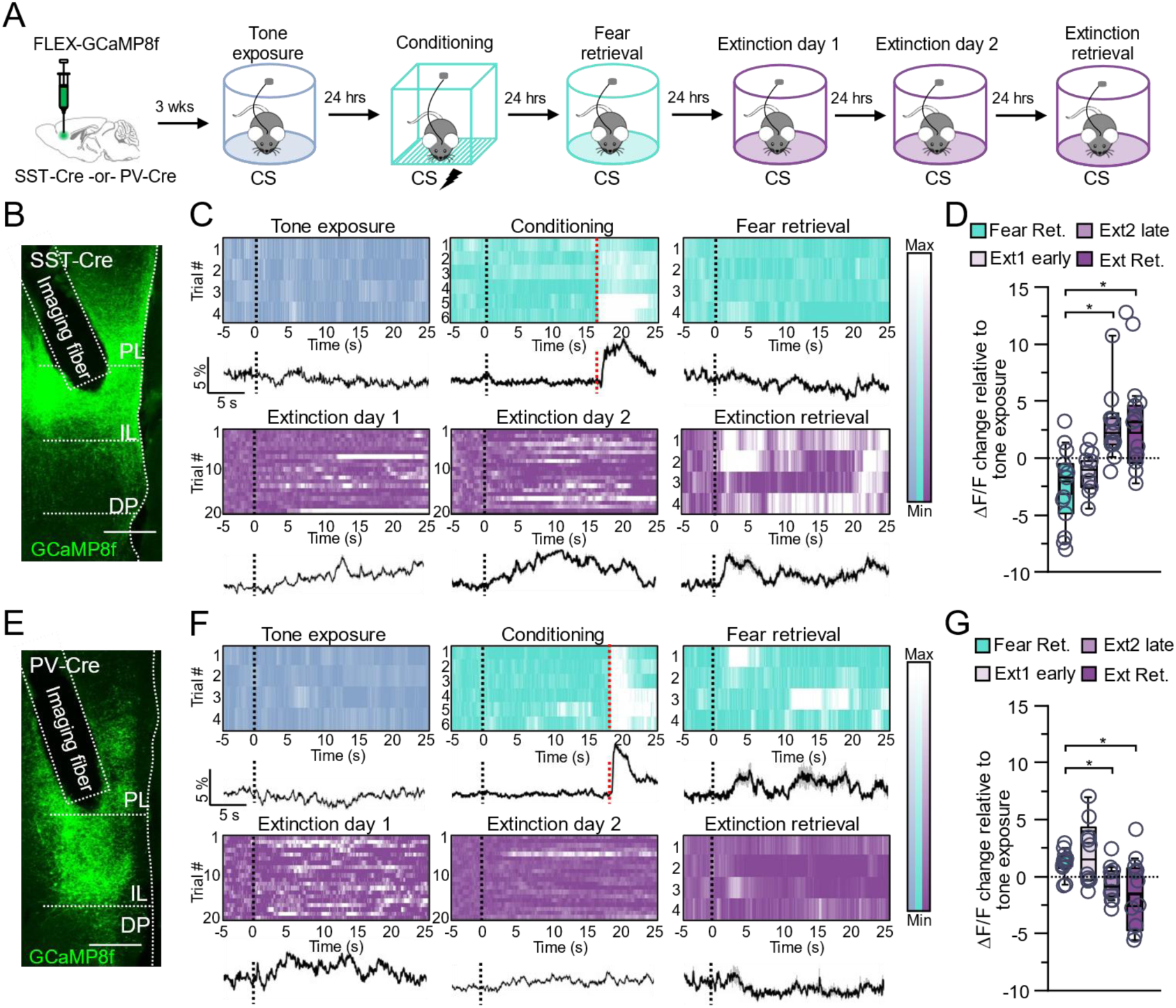
PV-INs and SST-INs exhibit dynamic orthogonal activity across fear and extinction behaviors. (A) SST-Cre and PV-Cre received infusions of FLEX-GCaMP8f in infralimbic cortex. An imaging fiber was also implanted immediately dorsal to infralimbic cortex. After 3 weeks, mice were subjected to a tone-exposure test consisting of four auditory tones (2 kHz, 80 db, 20 s). The next day, mice were then subjected to six pairings of auditory tones (2 kHz, 80 db, 20 s) and co-terminating foot shocks (0.7 mA, scrambled, 2 s). 24 hours later, mice were subjected to a fear memory retrieval test (4 CS, 2 kHz, 80 db, 20 s). Auditory fear memory extinction (20 CS tones per session, 2 kHz, 80 db, 20 s) was then performed for two consecutive days. Finally, mice were tested in a CS-evoked extinction memory retrieval task (2 kHz, 80 db, 20 s). Calcium signals were continuously recorded throughout all behavior testing. (B) Representative histological image of mPFC from SST-Cre mice expressing GCaMP8f and implanted imaging fiber. Scale = 300 µm. (C) Representative average calcium-associated fluorescence traces and associated heat maps for signals recorded from infralimbic cortex SST-INs during all behaviors. Black dotted lines represent CS onset and red dotted lines represent shock onset. Traces: black lines represent mean signal; gray represents SEM. Heat maps: dark color = minimum signal; white = maximum signal. (D) Changes in calcium signal across behavioral sessions compared to tone exposure (Χ^2^ = 8.71 [3], p = 0.01, Friedman ANOVA; n = 13 mice). (E) Representative histological image of mPFC from PV-Cre mice expressing GCaMP8f and implanted imaging fiber. Scale = 300 µm. (F) Representative average calcium-associated fluorescence traces and associated heat maps for signals recorded from infralimbic cortex PV-INs during all behaviors. Dotted lines represent CS onset for each trial. Heat maps: dark color = minimum signal; white = maximum signal. (G) Changes in calcium signal across behavioral sessions compared to tone exposure (Χ^2^ = 11.7 [3], p = 0.008, Friedman ANOVA; n = 12 mice). *, p < 0.05, Dunn’s post-hoc test. Boxplots represent the median (center line), mean (square), quartiles, and 10%–90% range (whiskers/error bars). Open circles represent data points for individual mice.

If IL SST-INs and PV-INs are differentially engaged across low and high fear states, respectively, we reasoned that manipulating their activity during extinction training should impact extinction encoding and retrieval in distinct ways. We therefore performed *in vivo* optogenetic activation or silencing experiments. Flp- or Cre-dependent channelrhodopsin (ChR2 for SST-INs; ChETA for PV-INs), archaerhodopsin (Arch), or control eYFP were bilaterally infused into IL of SST-Flp or PV-Cre mice (**Fig. 4a-c, 4n-o**). Optic fibers were also bilaterally implanted immediately above IL (**Supplementary Fig. 6**). After fear conditioning (**Supplementary Fig. 7**), mice were subjected to two consecutive days of extinction training during which IL SST-INs or PV-INs were optogenetically activated or silenced during each CS presentation. 24 hours later, mice were then subjected to extinction memory retrieval in the absence of optogenetic manipulation (**Fig. 4a**). While silencing IL SST-INs on extinction day 1 had no obvious effect on freezing (**Fig. 4d**), activating them reduced freezing to levels indistinguishable from baseline by the end of the training session (**Fig. 4e**). Normalization of CS-evoked freezing to baseline freezing during early (CS1-4) and late (CS17-20) periods of extinction 1 across all groups revealed that activation of IL SST-INs significantly facilitated extinction encoding (**Fig. 4f**). Performing the same manipulations during extinction day 2 resulted in more pronounced effects. Specifically, silencing IL SST-INs resulted in elevated freezing even in the late CS epochs, during which eYFP controls had fully extinguished (**Fig. 4g**). On the other hand, activating IL SST-INs reduced freezing to baseline-like levels during both early and late CS epochs, where eYFP controls were still freezing at levels significantly greater than baseline (**Fig. 4h**). Normalization of CS-evoked freezing to baseline freezing during early and late periods of extinction 2 across groups revealed that silencing IL SST-INs blocked extinction during both early and late periods, whereas activating these cells facilitated extinction encoding (**Fig. 4i**). Extinction memory retrieval was then performed in the absence of IL SST-IN manipulation. In mice that received IL SST-IN silencing during extinction training, we observed a significant increase in CS-evoked freezing over baseline levels that was absent in controls (**Fig. 4j**). Mice that received IL SST-IN activation during extinction training froze at comparable levels during both baseline and CS presentation epochs, similar to eYFP control, likely due to a floor effect (**Fig. 4k**). Normalization of CS-evoked freezing to baseline freezing revealed similar trends (**Fig. 4l**). These results suggest that IL SST-IN activity during extinction training facilitates extinction memory encoding and subsequent expression the next day (**Fig. 4m**).

**Figure 4.**
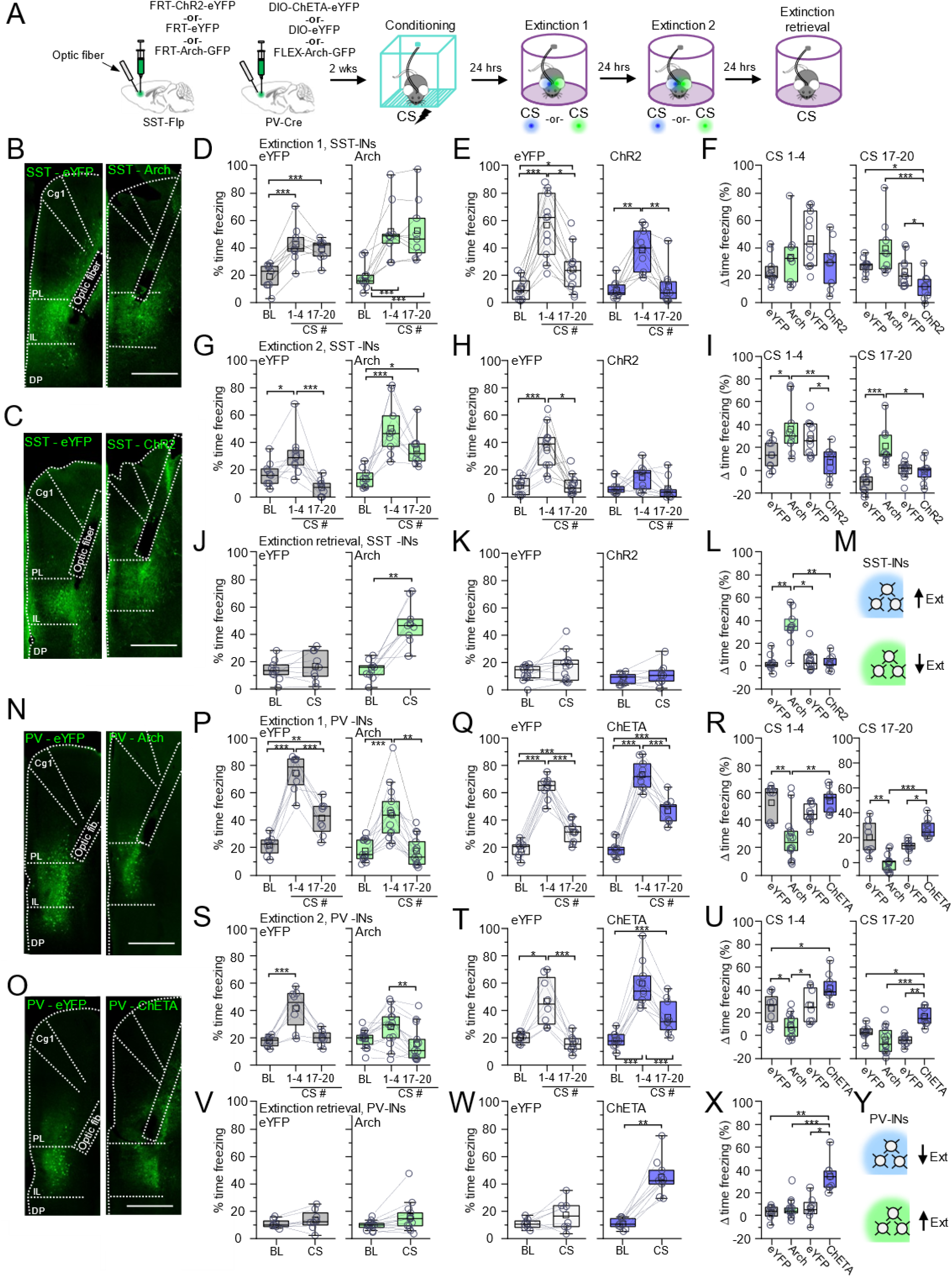
Infralimbic cortex SST-INs and PV-INs differentially drive fear and extinction behaviors. **(A)** SST-Flp mice received infusions of FRT-ChR2-eYFP (n = 9 mice), FRT-Arch-GFP (n = 9 mice), or FRT-eYFP (for ChR2 eYFP control: n = 11 mice; for Arch eYFP control: n = 9 mice) while PV-Cre mice received infusions DIO-ChETA-eYFP (n = 9 mice), FLEX-Arch-GFP (n = 13 mice), or DIO-eYFP (for ChETA eYFP control: n = 9 mice; for Arch eYFP control: n = 8 mice). Optical fibers were also implanted and directed at the infralimbic cortex. After 2 weeks, mice were subjected to six pairings of auditory tones (2 kHz, 80 db, 20 s) and co-terminating foot shocks (0.7 mA, scrambled, 2 s). 24 hours later, mice were subjected to auditory fear memory extinction (20 CS tones per session, 2 kHz, 80 db, 20 s) with simultaneous optogenetic activation or silencing during CS presentations for two consecutive days. Mice were then tested in a CS-evoked extinction memory retrieval task (2 kHz, 80 db, 20 s) without optogenetic manipulation. **(B)** Representative histological images of mPFC from SST-Flp mice expressing eYFP or Arch and implanted optic fibers. Scale = 500 µm. **(C)** Representative histological images of mPFC from SST-Flp mice expressing eYFP or ChR2 and implanted optic fibers. Scale = 500 µm. **(D)** Freezing quantified during CS (2 kHz, 20 s, 80 dB) and light (532 nm, solid light, 20 s epochs) presentation for animals expressing eYFP (F_2,16_ = 34.24, p = 0.0001, repeated-measures ANOVA) or Arch (F_2,16_ = 20.02, p = 0.0001, repeated-measures ANOVA) in infralimbic cortex SST-INs during early (CS 1-4) and late (CS 17-20) blocks of extinction day 1. **(E)** Freezing quantified during CS (2 kHz, 20 s, 80 dB) and light (473 nm, 20 Hz, 5 ms pulses, 20 s epochs) presentation for animals expressing eYFP (Χ^2^ = 22 [2], p = 0.0001, Friedman ANOVA) or ChR2 (Χ^2^ = 13.56 [2], p = 0.001, Friedman ANOVA) in infralimbic cortex SST-INs during early (CS 1-4) and late (CS 17-20) blocks of extinction day 1. **(F)** Quantification of CS-evoked freezing normalized to baseline freezing during early (CS 1-4; Χ^2^ = 7.28 [3], p = 0.06, Kruskal-Wallis) and late (CS 17-20; Χ^2^ = 15.94 [3], p = 0.001, Kruskal-Wallis) blocks of extinction day 1. **(G)** Freezing quantified during CS (2 kHz, 20 s, 80 dB) and light (532 nm, solid light, 20 s epochs) presentation for animals expressing eYFP (F_2,24_ = 10.43, p = 5.48×10^-4^, repeated-measures ANOVA) or Arch (Χ^2^ = 14.89 [2], p = 5.85×10^-4^, Friedman ANOVA) in infralimbic cortex SST-INs during early (CS 1-4) and late (CS 17-20) blocks of extinction day 2. **(H)** Freezing quantified during CS (2 kHz, 20 s, 80 dB) and light (473 nm, 20 Hz, 5 ms pulses, 20 s epochs) presentation for animals expressing eYFP (Χ^2^ = 14.36 [2], p = 7.6×10^-4^, Friedman ANOVA) or ChR2 (Χ^2^ = 2.67 [2], p = 0.26, Friedman ANOVA) in infralimbic cortex SST-INs during early (CS 1-4) and late (CS 17-20) blocks of extinction day 2. **(I)** Quantification of CS-evoked freezing normalized to baseline freezing during early (CS 1-4; F_3,34_ = 5.88, p = 0.002, one-way ANOVA) and late (CS 17-20; Χ^2^ = 20.59 [3], p = 1.28×10^-4^, Kruskal-Wallis) blocks of extinction day 2. **(J)** Freezing quantified during CS-evoked fear extinction memory retrieval (2 kHz, 20 s, 80 dB) for animals expressing eYFP (t_8_ = -1.2, p = 0.26, paired t-test) or Arch (W = 0, p = 0.009, Wilcoxon signed ranks test) in infralimbic cortex SST-INs. **(K)** Freezing quantified during CS-evoked fear extinction memory retrieval (2 kHz, 20 s, 80 dB) for animals expressing eYFP (W = 15, p = 0.12, Wilcoxon signed ranks test) or ChR2 (t_8_ = - 1.44, p = 0.19, paired t-test) in infralimbic cortex SST-INs. **(L)** Quantification of CS-evoked freezing normalized to baseline freezing during fear extinction memory retrieval (Χ^2^ = 16.71 [3], p = 8.11×10^-4^, Kruskal-Wallis) **(M)** Cartoon summary of findings indicating that optogenetic enhancement or suppression of infralimbic cortex SST-INs promotes and suppresses fear memory extinction, respectively. **(N)** Representative histological images of mPFC from PV-Cre mice expressing eYFP or Arch and implanted optic fibers. Scale = 500 µm. **(O)** Representative histological images of mPFC from PV-Cre mice expressing eYFP or ChETA and implanted optic fibers. Scale = 500 µm. **(P)** Freezing quantified during CS (2 kHz, 20 s, 80 dB) and light (532 nm, solid light, 20 s epochs) presentation for animals expressing eYFP (F_2,14_ = 57.22, p = 0.0001, one-way repeated-measures ANOVA) or Arch (Χ^2^ = 19.85 [2], p = 0.0001, Friedman ANOVA) in infralimbic cortex PV-INs during early (CS 1-4) and late (CS 17-20) blocks of extinction day 1. **(Q)** Freezing quantified during CS (2 kHz, 20 s, 80 dB) and light (473 nm, 50 Hz, 2 ms pulses, 20 s epochs) presentation for animals expressing eYFP (F_2,16_ = 176.3, p = 0.0001, one-way repeated-measured ANOVA) or ChETA (F_2,16_ = 163.5, p = 0.0001, one-way repeated-measures ANOVA) in infralimbic cortex PV-INs during early (CS 1-4) and late (CS 17-20) blocks of extinction day 1. **(R)** Quantification of CS-evoked freezing normalized to baseline freezing during early (CS 1-4; Χ^2^ = 17.61 [3], p = 5.29×10^-4^, Kruskal-Wallis) and late (CS 17-20; Χ^2^ = 25.33 [3], p = 0.0001, Kruskal-Wallis) blocks of extinction day 1. **(S)** Freezing quantified during CS (2 kHz, 20 s, 80 dB) and light (532 nm, solid light, 20 s epochs) presentation for animals expressing eYFP (Χ^2^ = 14.25 [2], p = 8.05×10^-4^, Friedman ANOVA) or Arch (Χ^2^ = 9.85 [2], p = 0.007, Friedman ANOVA) in infralimbic cortex PV-INs during early (CS 1-4) and late (CS 17-20) blocks of extinction day 2. **(T)** Freezing quantified during CS (2 kHz, 20 s, 80 dB) and light (473 nm, 50 Hz, 2 ms pulses, 20 s epochs) presentation for animals expressing eYFP (Χ^2^ = 14.25 [2], p = 8.05×10^-4^, Friedman ANOVA) or ChETA (F_2,16_ = 69.62, p = 0.0001, one-way repeated-measured ANOVA) in infralimbic cortex PV-INs during early (CS 1-4) and late (CS 17-20) blocks of extinction day 2. **(U)** Quantification of CS-evoked freezing normalized to baseline freezing during early (CS 1-4; F_3,34_ = 12.43, p = 0.0001, one-way ANOVA) and late (CS 17-20; Χ^2^ = 19.1 [3], p = 2.66×10^-4^, Kruskal-Wallis) blocks of extinction day 2. **(V)** Freezing quantified during CS-evoked fear extinction memory retrieval (2 kHz, 20 s, 80 dB) for animals expressing eYFP (W = 0, p = 0.23, Wilcoxon signed ranks test) or Arch (W = 11, p = 0.018, Wilcoxon signed ranks test) in infralimbic cortex PV-INs. **(W)** Freezing quantified during CS-evoked fear extinction memory retrieval (2 kHz, 20 s, 80 dB) for animals expressing eYFP (W = 6, p = 0.11, Wilcoxon signed ranks test) or ChETA (W = 0, p = 0.009, Wilcoxon signed ranks test) in infralimbic cortex PV-INs. **(X)** Quantification of CS-evoked freezing normalized to baseline freezing during fear extinction memory retrieval (Χ^2^ = 18.84 [3], p = 2.96×10^-4^, Kruskal-Wallis). **(Y)** Cartoon summary of findings indicating that optogenetic enhancement or suppression of infralimbic cortex PV-INs suppresses and promotes fear memory extinction, respectively. *, p < 0.05, **, p < 0.01, and ***, p < 0.001, Tukey’s or Dunn’s post-hoc test. Boxplots represent the median (center line), mean (square), quartiles, and 10%–90% range (whiskers/error bars). Open circles represent data points for individual mice.

We next performed similar optogenetic manipulations in IL PV-INs during extinction training (**Fig. 4n-o**). were not different from baseline (**Fig. 4p**). While activating IL PV-INs appeared to have a limited effect on freezing (**Fig. 4q**), normalization of freezing during early and late CS epochs to baseline freezing revealed that this manipulation suppressed extinction, whereas silencing enhances extinction compared to eYFP controls (**Fig. 4r**). Manipulations on extinction day 2 resulted in similar effects. While optogenetic silencing of IL PV-INs suppressed freezing (**Fig. 4s**), their activation enhanced freezing (**Fig. 4t**). Normalization revealed the same trends across groups, indicating that silencing IL PV-INs facilitates extinction whereas their activation suppresses extinction (**Fig. 4u**). During extinction memory retrieval the next day, no differences in baseline and CS-evoked freezing were noted in mice where IL PV-INs were silenced during extinction training (**Fig. 4v**). However, in mice where IL PV-INs were activated during extinction training, significant increases in CS-evoked freezing compared to baseline were observed (**Fig. 4w**). Normalization of CS freezing to baseline freezing during extinction memory retrieval confirmed that activation of IL PV-INs during extinction training enhances freezing (**Fig. 4x**). These results suggest IL PV-IN activity during extinction training suppresses extinction encoding and retrieval the next day (**Fig. 4y**).

Notably, activating or silencing IL SST-INs or PV-INs in a pre-conditioning tones exposure test does not impact freezing (**Supplemental Fig. 8**). Likewise, the same manipulations in the open field did not affect locomotion parameters (**Supplemental Fig. 9**). These control tests suggest that IL PV-IN and SST-IN activity is likely specifically related to fear and extinction memory processing, rather than signaling freezing or locomotion generally. These results are also in line with control fiber photometry analyses revealing IL SST-IN and PV-IN activity is not tied to freezing behaviors or locomotion occurring outside the CS (**Supplementary Fig. 5**). Taken together, these results suggest that IL SST-IN and PV-IN activity likely play opposite roles in extinction memory processes, where SST-IN and PV-IN activity enhances and suppresses extinction encoding, respectively.

Learning-related plasticity is thought to be an important mechanism for memory encoding^34,35^. Thus, distinct learning-related plasticity of IL SST-INs and PV-INs in response to fear conditioning and fear memory extinction may represent a potential mechanism for the differential recruitment of these cell types. We therefore recorded miniature excitatory (mEPSC) and inhibitory (mIPSC) postsynaptic currents from IL SST-INs and PV-INs. SST-Flp/ Ai65F and PV-Cre/ Ai9 mice were either home cage experienced (naïve group), or were subjected to fear conditioning only or to fear conditioning followed by two consecutive days of extinction training separated by 24 hours (extinction group) (**Fig 5a, Supplementary Fig. 10**). The next day, mice were used to prepare acute brain slices and IL SST-INs and PV-INs were targeted using red fluorescence (**Fig. 5b,g**). IL SST-INs recorded from mice in the extinction group exhibited increased frequency of mEPSCs compared to fear or naïve groups, with no group differences in amplitude (**Fig. 5c, 5d**). mIPSC recordings from the same IL SST-INs revealed no significant group differences in either frequency or amplitude (**Fig. 5e, 5f**). The same recordings were performed in IL PV-INs (**Fig. 5g**). IL PV-INs recorded from mice in the fear group exhibited increased mEPSC frequency compared to naïve and extinction groups with no group differences in amplitude (**Fig. 5h, 5i**). mIPSC recordings from the same IL PV-INs revealed increased frequency in cells recorded from the extinction group compared to fear and naïve groups with no group differences in amplitude (**Fig. 5j, 5k**). These results reveal that IL SST-INs receive enhanced excitatory drive following fear extinction. In contrast, IL PV-INs receive enhanced excitatory drive after fear conditioning which is lost after extinction. At the same time, IL PV-INs also receive enhanced inhibitory drive after fear extinction.

**Figure 5.**
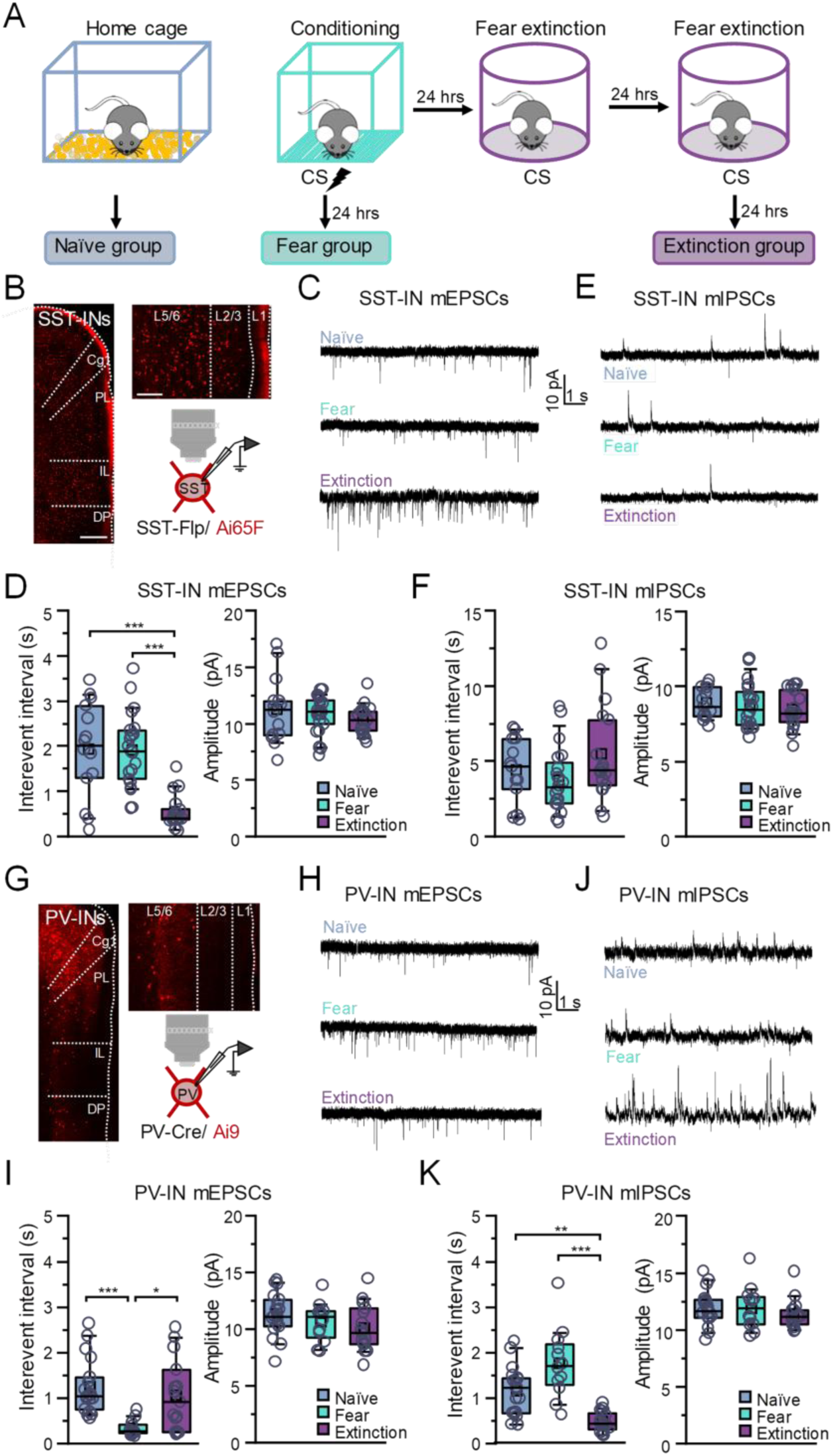
Infralimbic cortex PV-INs and SST-INs receive increased excitatory transmission after fear conditioning and extinction training, respectively. (A) Mice were experienced to their home cage and used for recordings (naïve group) or were subjected to six pairings of auditory tones (2 kHz, 80 db, 20 s) and co-terminating foot shocks (0.7 mA, scrambled, 2 s) and used for electrophysiological recordings 24 hours later (fear group) or to six pairings of auditory tones (2 kHz, 80 db, 20 s) and co-terminating foot shocks (0.7 mA, scrambled, 2 s) followed by two consecutive days of auditory fear memory extinction training (20 tones per session, 2 kHz, 80 db, 20 s) and used for electrophysiological recordings 24 hours later (extinction group). (B) Representative histological image from mPFC of SST-Flp/ Ai65F and cartoon of the recording configuration. (C) Representative miniature excitatory postsynaptic current (EPSC) recordings in infralimbic cortex SST-INs from brain slices prepared from naive (8 slices from 4 mice), fear conditioned (8 slices from 4 mice), and extinction (12 slices from 6 mice) mice. (D) Quantification of the mEPSC interevent interval and amplitude for SST-INs in naive, fear, and extinction groups. Interevent interval: Χ^2^ = 23.91 (2), p = 0.0001, Kruskal-Wallis; naive = 14 cells; fear = 21 cells; extinction = 18 cells. Amplitude: Χ^2^ = 1.59 (2), p = 0.45, Kruskal-Wallis; naive = 14 cells; fear = 21 cells; extinction = 18 cells. (E) Representative miniature inhibitory postsynaptic current (mIPSC) recordings in infralimbic cortex SST-INs from brain slices prepared from naive (8 slices from 4 mice), fear conditioned (8 slices from 4 mice), and extinction (12 slices from 6 mice) mice. (F) Quantification of the mIPSC interevent interval and amplitude for naive, fear, and extinction groups. Interevent interval: F_2,47_ = 2.06, p = 0.14, one-way ANOVA; naive = 14 cells; fear = 21 cells; extinction = 18 cells. Amplitude: F_2,47_ = 0.46, p = 0.64, one-way ANOVA; naive = 14 cells; fear = 21 cells; extinction = 18 cells. (G) Representative histological image from mPFC of PV-Cre/ Ai9 and cartoon of the recording configuration. (H) Representative miniature excitatory postsynaptic current (mEPSC) recordings in infralimbic cortex PV-INs from brain slices prepared from naive (6 slices from 3 mice), fear conditioned (8 slices from 4 mice), and extinction (8 slices from 4 mice) mice. (I) Quantification of the mEPSC interevent interval and amplitude for PV-INs in naive, fear, and extinction groups. Interevent interval: Χ^2^ = 17.32 (2), p = 1.73×10^-4^, Kruskal-Wallis; naive = 18 cells; fear = 12 cells; extinction = 15 cells. Amplitude: F_2,42_ = 1.61, p = 0.21, one-way ANOVA; naive = 18 cells; fear = 12 cells; extinction = 15 cells. (J) Representative miniature inhibitory postsynaptic current (mIPSC) recordings in infralimbic cortex PV-INs from brain slices prepared from naive (6 slices from 3 mice), fear conditioned (8 slices from 4 mice), and extinction (8 slices from 4 mice) mice. (K) Quantification of the mIPSC interevent interval and amplitude for naive, fear, and extinction groups. Interevent interval: Χ^2^ = 24.67 (2), p = 0.0001, Kruskal-Wallis; naive = 18 cells; fear = 12 cells; extinction = 15 cells. Amplitude: Χ^2^ = 1.36 (2), p = 0.51, Kruskal-Wallis; naive = 18 cells; fear = 12 cells; extinction = 15 cells. *, p < 0.05, **, p < 0.01, and ***, p < 0.001, Tukey’s or Dunn’s post-hoc test. Boxplots represent the median (center line), mean (square), quartiles, and 10%–90% range (whiskers/error bars). Open circles represent data points for individual cells.

The increased mIPSC drive onto PV-INs in the extinction group (**Fig. 5j,k**) may reflect an important disinhibitory mechanism for the recruitment of IL PNs and thus for extinction encoding. Since our data suggest IL SST-INs are recruited by extinction processes, this source of inhibition onto PV-INs may originate with IL SST-INs. To test this possibility, we performed optogenetics-assisted IL microcircuit electrophysiological analyses. Flp-dependent or Cre-dependent ChR2 was infused into SST-Flp/ PV-Cre/ Ai3 or SST-Flp/ PV-Cre/ Ai65F triple transgenic mice, respectively (**Fig. 6a**). Mice were split into three groups: naïve, fear, and extinction groups (**Supplementary Fig. 11**). The next day, we prepared brain slices from these mice. Using these vectors and mouse lines, it is possible to perform light-elicited responses from one IL interneuron population (e.g., PV-INs) onto another IL interneuron population identified with fluorescence (e.g., SST-INs) and neighboring IL PNs in the same slices (**Fig. 6b**). First, we recorded light-elicited responses from IL SST-INs onto PV-INs and PNs (**Fig. 6c**). Compared to naïve and fear groups, where SST-IN input was seemingly more biased onto PNs than PV-INs, the same recordings in the extinction group revealed SST-IN input onto these cells was roughly similar (**Fig. 6d-e**). Normalization of SST-IN input onto PV-INs to PNs in the same slices revealed that fear extinction training results in an increase in the PV:PN ratio compared to naïve or fear groups (**Fig. 6f**). Light-elicited responses from IL PV-INs onto SST-INs and PNs were also performed (**Fig. 6g**). Regardless of group, the balance of inhibition from IL PV-INs onto SST-INs and PNs was not different (**Fig. 6h-j**). To test whether the apparent change in SST-IN inhibition onto PV-INs after extinction training is due to learning-related alterations in GABA release, paired pulse recordings were performed. In line with this prediction, we observed a decrease in the inhibitory paired-pulse ratio in the extinction group compared to naïve and fear groups (**Fig. 6k-l**). The same paired-pulse recordings were performed for PV-IN inhibition onto SST-INs. Consistent with the idea that this synaptic connection likely does not change across groups, we did not observe group-dependent differences in paired-pulse ratio (**Fig. 6m-n**). Overall, these results suggest that synaptic connections from IL SST-INs onto PV-INs are specifically potentiated after extinction training (**Fig. 6o-p**), likely reflecting a disinhibitory circuit mechanism in which IL PNs are subsequently recruited.

**Figure 6.**
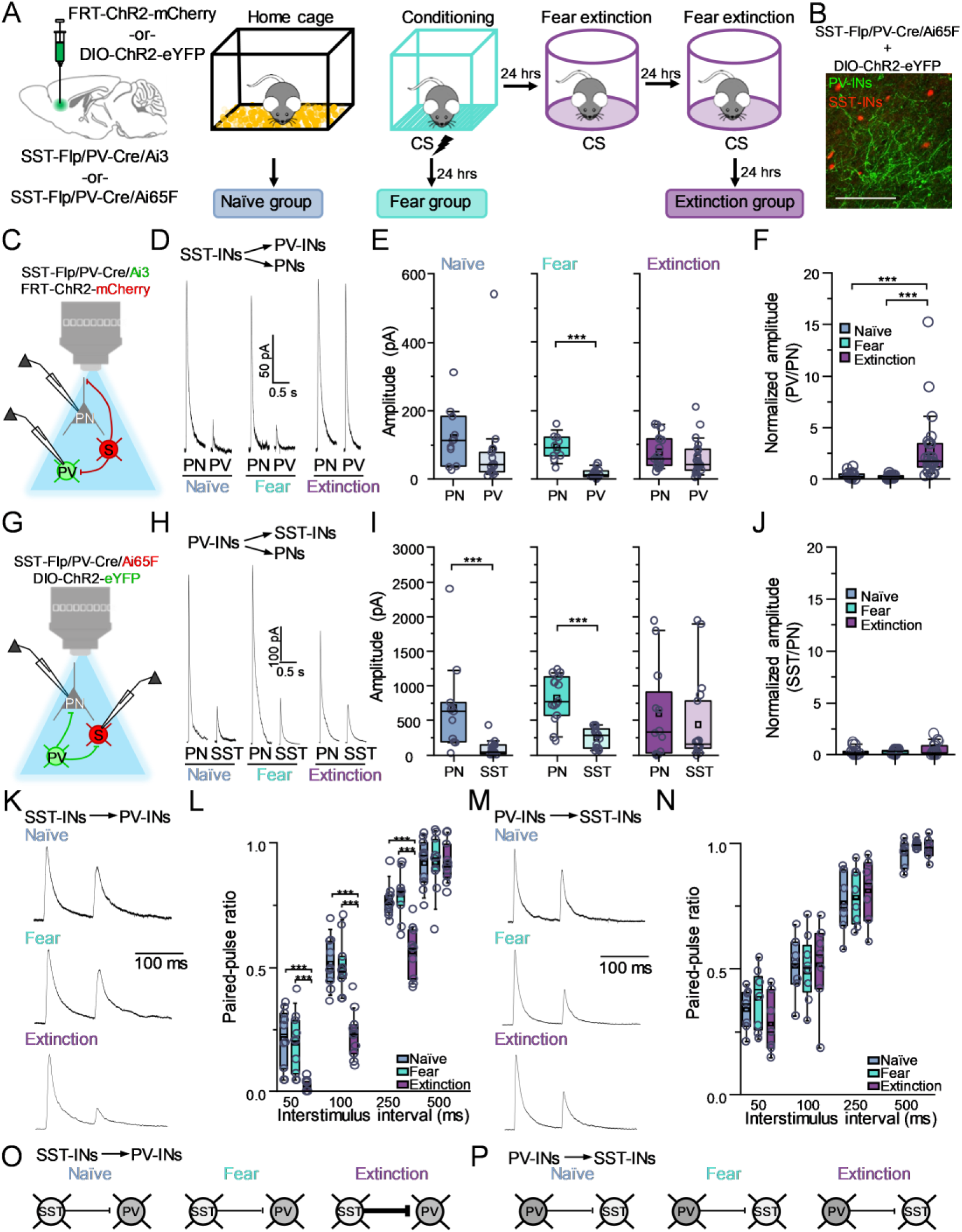
Infralimbic cortex SST-INs preferentially inhibit PV-INs after fear extinction training. (A) SST-Flp/ PV-Cre/ Ai3 and SST-Flp/ PV-Cre/ Ai65F triple transgenic mice received infusion of FRT-ChR2-mCherry and DIO-ChR2-eYFP, respectively, in the infralimbic cortex. After two weeks, mice were randomly split into three groups. Mice were either experienced to their home cage and used for recordings (naïve group); subjected to six pairings of auditory tones (2 kHz, 80 db, 20 s) and co-terminating foot shocks (0.7 mA, scrambled, 2 s) and used for electrophysiological recordings 24 hours later (fear group); or subjected to six pairings of auditory tones (2 kHz, 80 db, 20 s) and co-terminating foot shocks (0.7 mA, scrambled, 2 s) followed by two consecutive days of fear memory extinction training (20 tones per session, 2 kHz, 80 db, 20 s) and used for electrophysiological recordings 24 hours later (extinction group). (B) Representative histological image of DIO-ChR2-eYFP infused in infralimbic cortex of SST-Flp/ PV-Cre/ Ai65F. Scale = 100 µm. (C) Cartoon representation of whole-cell recording configuration for SST-Flp/ PV-Cre/ Ai3 mice infused with FRT-ChR2-mCherry. (D) Representative light-evoked inhibitory postsynaptic current (IPSC) recordings from infralimbic cortex SST-INs onto neighboring PV-INs and PNs in mice from naïve, fear, and extinction groups. (E) Quantification of light-evoked IPSCs from infralimbic cortex SST-INs onto neighboring PV-INs and PNs for naïve (U = 104, p = 0.06, Mann-Whitney U-test; n = 11 PNs and 13 PV-INs from 6 slices from 4 mice), fear (U = 188, p = 0.0001, Mann-Whitney U-test; n = 10 PNs and 19 PV-INs from 7 slices from 4 mice), and extinction (U = 239, p = 0.16, Mann-Whitney U-test; n = 18 PNs and 21 PV-INs from 8 slices from 4 mice) groups. (F) Light-evoked IPSCs from infralimbic cortex SST-INs onto neighboring PV-INs normalized to PNs from the same slices (Χ^2^ = 35.2 [2], p = 0.0001, Kruskal-Wallis). (G) Cartoon representation of whole-cell recording configuration for SST-Flp/ PV-Cre/ Ai65F mice infused with DIO-ChR2-eYFP. (H) Representative light-evoked inhibitory postsynaptic current (IPSC) recordings from infralimbic cortex PV-INs onto neighboring SST-INs and PNs in mice from naïve, fear, and extinction groups. (I) Quantification of light-evoked IPSCs from infralimbic cortex PV-INs onto neighboring SST-INs and PNs for naïve (U = 149, p = 6.14×10^-4^, Mann-Whitney U-test; n = 11 PNs and 15 SST-INs from 7 slices from 4 mice), fear (U = 249, p = 0.0001, Mann-Whitney U-test; n = 15 PNs and 18 SST-INs from 5 slices from 3 mice), and extinction (U = 121, p = 0.61, Mann-Whitney U-test; n = 12 PNs and 18 SST-INs from 8 slices from 4 mice) groups. (J) Light-evoked IPSCs from infralimbic cortex PV-INs onto neighboring SST-INs normalized to PNs from the same slices (Χ^2^ = 1.73 [2], p = 0.42, Kruskal-Wallis). (K) Representative light-evoked IPSC recordings from infralimbic cortex SST-INs onto PV-INs during paired pulse stimulation. (L) Quantification of paired-pulse ratio (F_6,54_ = 6.52, p = 0.0001, interaction between training and delay, two-way repeated-measures ANOVA; naïve: n = 10 PV-INs from 5 slices from 4 mice; fear: n = 10 PV-INs from 4 slices from 4 mice; n = 11 PV-INs from 5 slices from 4 mice). (M) Representative light-evoked IPSC recordings from infralimbic cortex PV-INs onto SST-INs during paired pulse stimulation. (N) Quantification of paired-pulse ratio (F_6,48_ = 1.76, p = 0.304, two-way repeated-measures ANOVA; naïve: n = 7 SST-INs from 3 slices from 3 mice; fear: n = 8 SST-INs from 3 slices from 3 mice; n = 7 SST-INs from 3 slices from 3 mice). (O) Cartoon summary of infralimbic cortex microcircuit balance alterations for SST-IN input onto PV-INs across naïve, fear, and extinction groups. (P) Cartoon summary of infralimbic cortex microcircuit balance alterations for PV-IN input onto SST-IN across naïve, fear, and extinction groups. ***, p < 0.001, Tukey’s post-hoc test. Boxplots represent the median (center line), mean (square), quartiles, and 10%–90% range (whiskers/error bars). Open circles are data points for individual cells.

The BLA is a locus for both fear encoding and fear memory extinction^36–39^, and sends dense projections to IL^40^ that are important for fear extinction^39,41^. We therefore tested whether BLA might drive shifts in enhanced excitation onto IL PV-INs versus IL SST-INs after fear conditioning and fear memory extinction, respectively, as observed previously (**Fig. 5**). SST-Flp/ Ai65F and PV-Cre/ Ai9 mice received bilateral infusions of CaMKII-ChR2-eYFP in BLA and split into three groups: naïve, fear, and extinction groups (**Fig. 7a; Supplementary Fig. 12**). 24 hours after the last day of training, prefrontal brain sections were procured and light-elicited responses from BLA terminals onto IL SST-INs, PV-INs, and PNs were recorded (**Fig. 7b-d**). In recordings from IL SST-INs and PNs, excitatory drive from BLA was seemingly weaker onto SST-INs compared to PNs in the naïve and fear groups compared to the extinction group (**Fig. 7e-f**). Normalization of responses onto IL SST-INs to PNs from the same slices revealed an apparent increase in excitatory drive from BLA onto SST-INs after extinction compared to naïve and fear groups (**Fig. 7g**). The same recordings onto IL PV-INs and PNs revealed an apparent shift in higher excitatory drive onto PV-INs versus PNs in the fear group compared to either naïve or extinction groups (**Fig. 7h-i**). Normalization of BLA responses onto PV-INs and PNs in the same slices revealed the same effects across groups (**Fig. 7j**). To test whether the apparent changes in excitatory synaptic drive might be due to experience-dependent presynaptic plasticity, paired-pulse recordings were performed. Paired pulse ratio across shorter delays was lower in IL SST-INs recorded in the extinction group compared to naïve or fear groups (**Fig. 7k-l**). On the other hand, paired-pulse ratio was lower in IL PV-INs in the fear group compared to naïve and extinction groups (**Fig. 7m-n**). Together, these results suggest that BLA excitatory drive onto IL SST-INs is enhanced after extinction (**Fig. 7o**), whereas BLA excitatory drive onto IL PV-INs is enhanced after conditioning (**Fig. 7p**). Overall, this dynamic shift in BLA excitatory drive onto IL SST-INs and PV-INs may represent an important mechanism for driving fear extinction encoding in the IL.

**Figure 7.**
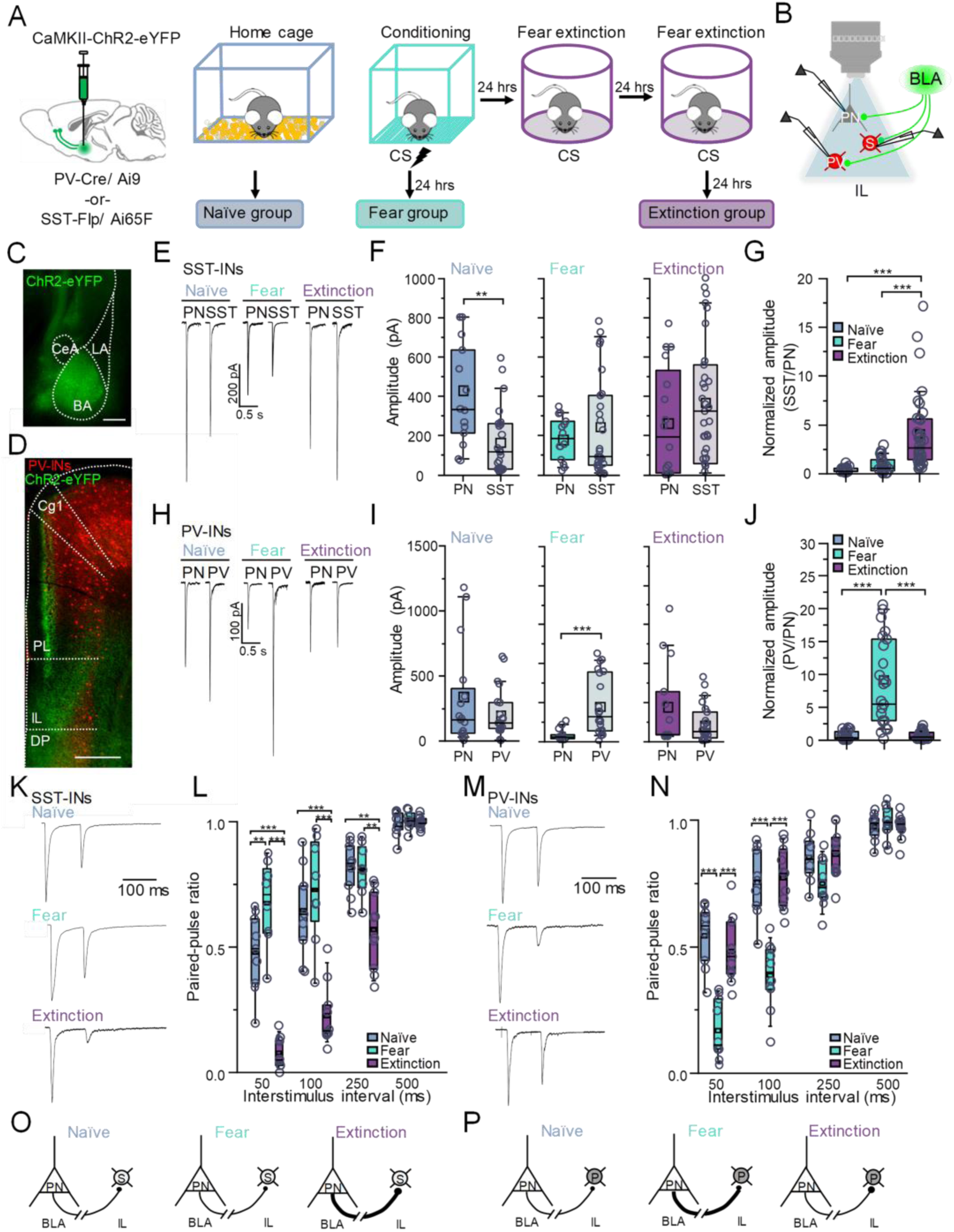
Relative balance of BLA inputs onto infralimbic cortex SST-INs and PV-INs are altered across fear and extinction training. (A) SST-Flp/ Ai65F or PV-Cre/ Ai9 mice received bilateral infusions of CaMKII-ChR2-eYFP in BLA. After 2 weeks, mice were randomly split into three groups: those that were experienced to their home cage and used for recordings (naïve group); those that were subjected to six pairings of auditory tones (2 kHz, 80 db, 20 s) and co-terminating foot shocks (0.7 mA, scrambled, 2 s) and used for electrophysiological recordings 24 hours later (fear group); or those that were subjected to six pairings of auditory tones (2 kHz, 80 db, 20 s) and co-terminating foot shocks (0.7 mA, scrambled, 2 s) followed by two days of fear memory extinction training (20 tones, 2 kHz, 80 db, 20 s) and used for electrophysiological recordings 24 hours later (extinction group). (B) Cartoon representation of whole-cell recording configuration. (C) Histological image of ChR2-eYFP expression in BLA. Scale = 50 µm. (D) Histological image of mPFC from a PV-Cre/ Ai9 mouse expressing ChR2-eYFP in BLA projections. Scale = 500 µm. (E) Representative light-evoked excitatory postsynaptic current (EPSC) recordings from BLA terminals onto infralimbic cortex SST-INs and PNs in mice from naïve, fear, and extinction groups. (F) Quantification of light-evoked EPSCs from BLA terminals onto infralimbic cortex SST-INs and PNs for naïve (U = 257, p = 0.003, Mann-Whitney U-test; n = 14 PNs and 23 SST-INs from 10 slices from 5 mice), fear (U = 220, p = 0.81, Mann-Whitney U-test; n = 14 PNs and 30 SST-INs from 9 slices from 5 mice), and extinction (U = 204, p = 0.329, Mann-Whitney U-test; n = 16 PNs and 31 SST-INs from 10 slices from 5 mice) groups. (G) Light-evoked EPSCs from BLA terminals onto infralimbic cortex SST-INs normalized to PNs from the same slices (Χ^2^ = 44.2 [2], p = 0.0001, Kruskal-Wallis). (H) Representative light-evoked excitatory postsynaptic current (EPSC) recordings from BLA terminals onto infralimbic cortex PV-INs and PNs in mice from naïve, fear, and extinction groups. (I) Quantification of light-evoked EPSCs from BLA terminals onto infralimbic cortex PV-INs and PNs for naïve (U = 181, p = 0.70, Mann-Whitney U-test; n = 16 PNs and 21 PV-INs from 10 slices from 5 mice), fear (U = 34, p = 4.34×10^-4^, Mann-Whitney U-test; n = 13 PNs and 20 PV-INs from 9 slices from 5 mice), and extinction (U = 172, p = 0.47, Mann-Whitney U-test; n = 13 PNs and 23 PV-INs from 9 slices from 5 mice) groups. (J) Light-evoked EPSCs from BLA terminals onto infralimbic cortex PV-INs normalized to PNs from the same slices (Χ^2^ = 32.3 [2], p = 0.0001, Kruskal-Wallis). (K) Representative light-evoked EPSC recordings from BLA terminals onto infralimbic cortex SST-INs during paired pulse stimulation. (L) Quantification of paired-pulse ratio (F_6,42_ = 13.02, p = 0.0001, interaction between training and delay, two-way repeated-measures ANOVA; naïve: n = 9 SST-INs from 3 slices from 3 mice; fear: n = 8 SST-INs from 3 slices from 3 mice; n = 10 SST-INs from 3 slices from 3 mice). (M) Representative light-evoked EPSC recordings from BLA terminals onto infralimbic cortex PV-INs during paired pulse stimulation. (N) Quantification of paired-pulse ratio (F_6,48_ = 9.83, p = 0.0001, interaction between training and delay, two-way repeated-measures ANOVA; naïve: n = 9 PV-INs from 3 slices from 3 mice; fear: n = 10 PV-INs from 3 slices from 3 mice; n = 11 PV-INs from 3 slices from 3 mice). (O) Cartoon summary of the balance of BLA inputs onto infralimbic cortex SST-INs across naïve, fear, and extinction groups. (P) Cartoon summary of the balance of BLA inputs onto infralimbic cortex PV-INs across naïve, fear, and extinction groups. *, p < 0.05, **, p < 0.01, ***, p < 0.001, Tukey’s or Dunn’s post-hoc test. Boxplots represent the median (center line), mean (square), quartiles, and 10%–90% range (whiskers/error bars). Open circles are data points for individual cells.

Finally, we tested the role for BLA projections to IL in driving extinction-related behavior and plasticity. We therefore performed *in vivo* optogenetic silencing of BLA projections to IL and subsequent electrophysiological recordings in the same mice. SST-Flp/ Ai65F or PV-Cre/ Ai9 mice received bilateral infusions of CaMKII-Arch-GFP or control CaMKII-eYFP and optic fiber implantation immediately above IL (**Supplemental Fig. 13**). Mice were subjected to conditioning in the absence of optogenetic silencing followed by two consecutive days of extinction training during which CS presentation was paired with optogenetic silencing of BLA terminals in IL. The next day, mice were either subjected to a fear extinction retrieval test or used to procure brain slices for whole-cell electrophysiological recordings (**Fig. 8a-b**). Behavioral analysis across extinction days 1 and 2 revealed that optogenetic silencing of BLA projections to IL significantly enhanced freezing. In line with this idea, mice expressing Arch exhibited significantly higher CS-evoked freezing compared to baseline, whereas eYFP control mice exhibited low freezing across both epochs, as expected for mice that properly extinguished the fear memory (**Fig. 8c**). Thus, these results suggest the activity of BLA to IL projections during extinction training is critical for fear extinction encoding. To test whether inhibition of these terminals during extinction training is sufficient to block extinction learning-related plasticity, we subsequently recorded mEPSCs and mIPSCs from IL SST-INs and PV-INs. In mice expressing Arch, we observed lower frequency of mEPSCs in IL SST-INs compared to the same recordings in mice expressing eYFP, with no group-dependent differences in amplitude (**Fig. 8d-e**). Neither mIPSC frequency nor amplitude were altered in IL SST-INs recorded from Arch-versus eYFP-expressing mice (**Fig. 8f-g**). For IL PV-INs, mEPSC frequency was higher in PV-INs recorded from Arch-compared to eYFP-expressing mice, with no difference in amplitude (**Fig. 8h-i**). Neither mIPSC frequency nor amplitude were altered in IL PV-INs recorded from Arch-versus eYFP-expressing mice (**Fig. 8j-k**). Thus, silencing BLA terminals in IL during extinction training specifically recapitulates behaviors and electrophysiological signatures consistent with the high fear state observed after conditioning, underscoring the critical contributions of these projections and their plasticity to the extinction of auditory fear memory.

**Figure 8.**
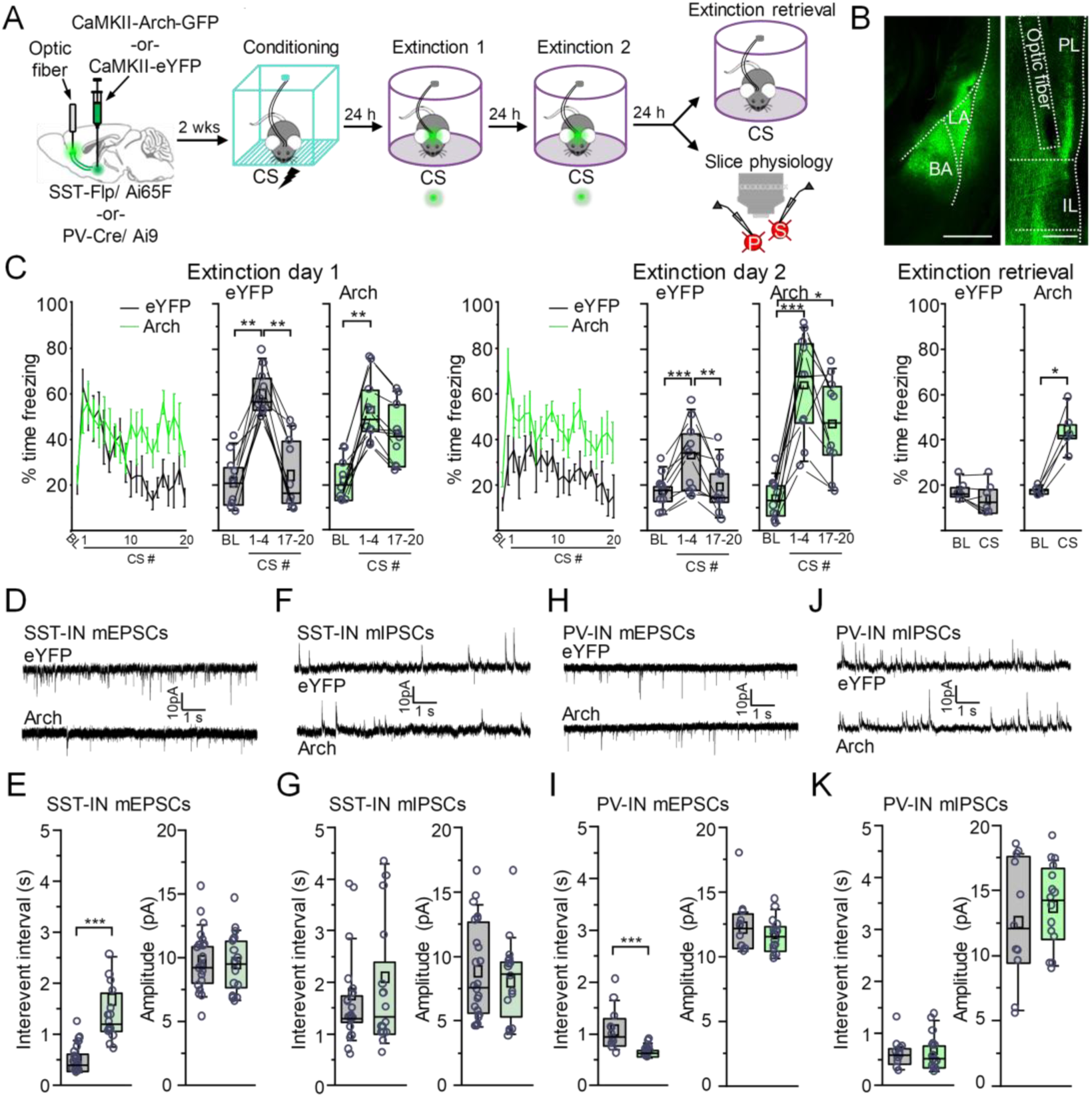
BLA projections to infralimbic cortex are required for extinction-related behavior and plasticity. **(A)** SST-Flp/ Ai65F and PV-Cre/ Ai9 mice received infusions of CaMKI-Arch or CaMKII-eYFP bilaterally in BLA. Optical fibers were implanted and directed at the infralimbic cortex. After 2 weeks, mice were subjected to six pairings of auditory tones (2 kHz, 80 db, 20 s) and co-terminating foot shocks (0.7 mA, scrambled, 2 s). 24 hours later, mice were subjected to two consecutive days of auditory fear memory extinction (20 CS tones per session, 2 kHz, 80 db, 20 s) with simultaneous optogenetic silencing during CS presentations. At 24 hours after combined training and optogenetic silencing, mice were used for whole-cell brain slice recordings or were subjected to an extinction memory retrieval test (4 CS tones, 2 kHz, 80 db, 20 s) in the absence of optogenetic silencing. **(B)** Representative histological images of CaMKII-Arch infusion in the basolateral amygdala (scale = 500 um) as well as the associated axon projections in mPFC (scale = 250 um). **(C)** Freezing quantified during CS (2 kHz, 20 s, 80 dB) and light (532 nm, solid light, 20 s epochs) presentation for animals expressing eYFP (Χ^2^ = 15 [2], p = 5.5 x 10^-4^, Friedman ANOVA; n = 10 mice) or Arch (Χ^2^ = 8.6 [2], p = 0.013, Friedman ANOVA; n = 10 mice) in BLA projections to infralimbic cortex during baseline and early (CS 1-4) and late (CS 17-20) blocks of extinction day 1; the same for the same animals expressing eYFP (F(2,18) = 13.1, p = 3.1 x 10^-4^, one-way repeated measures ANOVA) or Arch (Χ^2^ = 16.8 [2], p = 1.7 x 10^-4^, Friedman ANOVA) during baseline and early (CS 1-4) and late (CS 17-20) blocks of extinction day 2; and without optogenetic silencing for animals expressing eYFP (t5 = 1.86, p = 0.12, paired t-test) or Arch (W = 0, p = 0.041, Wilcoxon signed rank test) during baseline and CS presentation during extinction memory retrieval. **(D)** Representative miniature excitatory postsynaptic current (mEPSC) recordings in infralimbic cortex SST-INs from brain slices prepared from SST-Flp/ Ai65F mice expressing eYFP (8 slices from 4 mice) or Arch (8 slices from 4 mice) in BLA projections to infralimbic cortex. **(E)** Quantification of the mEPSC interevent interval and amplitude from infralimbic cortex SST-IN recordings for mice expressing eYFP or Arch in BLA projections to infralimbic cortex. Interevent interval: U = 67, p = 4.6×10^-4^, Mann-Whitney U-test; eYFP = 23 cells; Arch = 17 cells. Amplitude: t38 = 0.24; p = 0.81, two-sample t-test; eYFP = 23 cells; Arch = 17 cells. **(F)** Representative miniature inhibitory postsynaptic current (IPSC) recordings in infralimbic cortex SST-INs from brain slices prepared from SST-Flp/ Ai65F mice expressing eYFP (8 slices from 4 mice) or Arch (8 slices from 4 mice) in BLA projections to infralimbic cortex. **(G)** Quantification of the mIPSC interevent interval and amplitude for recordings from infralimbic cortex SST-INs in mice expressing eYFP or Arch in BLA projections to infralimbic cortex. Interevent interval: U = 173, p = 0.94, Mann-Whitney U-test; eYFP = 23 cells; Arch = 17 cells. Amplitude: t38 = 0.66; p = 0.52, two-sample t-test; eYFP = 23 cells; Arch = 17 cells. **(H)** Representative miniature excitatory postsynaptic current (mEPSC) recordings in infralimbic cortex PV-INs from brain slices prepared from PV-Cre/ Ai9 mice expressing eYFP (8 slices from 4 mice) or Arch (8 slices from 4 mice) in BLA projections to infralimbic cortex. **(I)** Quantification of the mEPSC interevent interval and amplitude in recordings from infralimbic cortex PV-INs in mice expressing eYFP or Arch in BLA projections to infralimbic cortex. Interevent interval: U = 149, p = 6.14×10^-4^; Amplitude: t24 = 1.14, p = 0.27, two-sample t-test; eYFP = 11 cells; Arch = 15 cells. **(J)** Representative miniature inhibitory postsynaptic current (mIPSC) recordings in infralimbic cortex PV-INs from brain slices prepared from PV-Cre/ Ai9 mice expressing eYFP (8 slices from 4 mice) or Arch (8 slices from 4 mice) in BLA projections to infralimbic cortex. **(K)** Quantification of the mIPSC interevent interval and amplitude for recordings from infralimbic cortex PV-INs for mice expressing eYFP or Arch in BLA projections to infralimbic cortex. Interevent interval: t24 = -0.16, p = 0.87; eYFP = 11 cells; Arch = 15 cells. Amplitude: t24 = -0.74, p = 0.46; eYFP = 11 cells; Arch = 15 cells. *, p < 0.05, **, p < 0.01, ***, p < 0.001, Tukey’s or Dunn’s post-hoc test. Boxplots represent the median (center line), mean (square), quartiles, and 10%–90% range (whiskers/error bars). Open circles represent data points for individual mice or cells.

## Discussion

In the current study, we revealed that the dynamic activity and plasticity of the IL GABAergic microcircuit is critical for the extinction of auditory fear memory (**Summary Figure**). While the role for IL in extinction behaviors has been known for decades, the exact cell and circuit plasticity mechanisms underlying this role remain underexplored. Using activity-dependent neural tagging and immunohistochemistry, we found that IL SST-INs and PV-INs are highly recruited following fear extinction and fear memory retrieval, respectively. *In vivo* calcium imaging further revealed that these cell populations exhibit orthogonal patterns of activity, where PV-INs were more highly recruited in high fear states like fear memory retrieval and early extinction epochs, whereas IL SST-INs were recruited during lower fear states that were learned, like later extinction epochs and extinction memory retrieval tests. *In vivo* optogenetic manipulations of IL SST-INs and PV-INs during extinction training revealed that the activity of SST-INs facilitates extinction, while the activity of PV-INs suppresses extinction by driving conditioned fear. Brain slice recordings revealed that IL SST-INs and PV-INs receive greater excitatory drive after extinction training and fear conditioning, respectively; PV-INs additionally receive greater inhibitory drive after extinction training. Optogenetics-assisted electrophysiological recordings revealed that IL SST-INs provide enhanced inhibition onto PV-INs, likely driving disinhibition of PNs through relief of PV-IN-mediated inhibition. At the same time, BLA dynamically engages the IL microcircuit, where SST-INs and PV-INs are preferentially engaged after fear extinction and fear conditioning, respectively. Silencing projections from BLA to IL during extinction training blocked fear extinction-related behavior and plasticity, underscoring the importance of this pathway in shaping IL microcircuit activity and extinction memory encoding. Taken together, these results reveal how the local circuits within IL are engaged to drive learned high or low fear behavioral states – a mechanism not only relevant for fundamental emotional memory processing, but for disorders where these circuits go awry such as PTSD^42,43^.

**Summary Figure.**
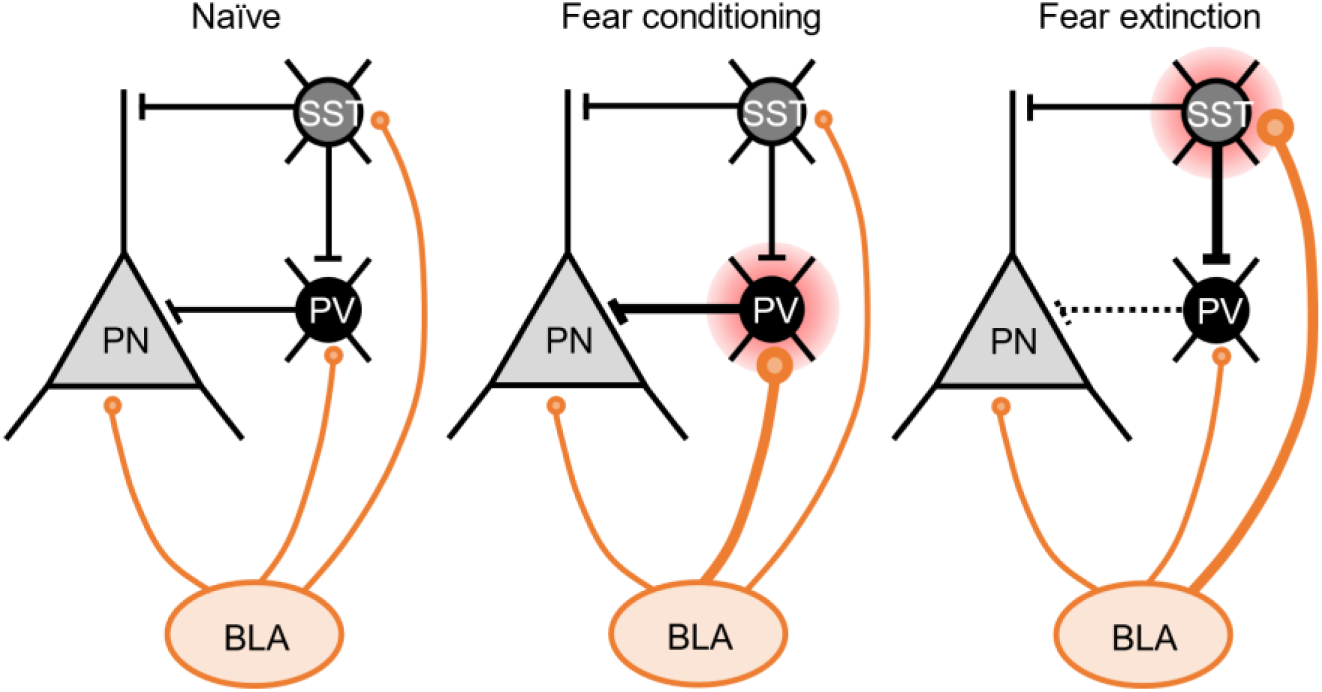
Following fear conditioning, BLA glutamatergic transmission is potentiated onto IL PV-INs, potentially driving their recruitment and feed-forward inhibition of IL PN activity. After fear extinction, the fear-related increase in BLA glutamatergic transmission onto IL PV-INs is reversed. At the same time, BLA glutamatergic transmission onto IL SST-INs is potentiated. In addition, fear extinction also drives enhanced inhibition from IL SST-INs specifically onto IL PV-INs, driving the disinhibition of IL principal neurons.

The role for IL in extinction has been known for decades. However, the local microcircuit computations that underlie this role had been largely unstudied. IL SST-INs in adolescent rodents have been implicated in the regulation of fear extinction, although the mechanisms we observe in adult rodents are distinct likely due to the differences in the developing circuit^29^. In adult rodents, stress-induced deficits in fear extinction are driven at least in part by enhanced activity of IL PV-INs^28^. In agreement with this finding, IL PV-IN network plasticity has been observed to correlate with successful fear extinction encoding^44^. We found that IL SST-INs and PV-INs have divergent roles in fear extinction processes. In agreement with Binette et al^28^, we find that the activity of IL PV-INs is linked to high fear states like fear memory retrieval. In addition, we also made the novel observation that IL SST-IN activity is enhanced during cued fear memory extinction. This new piece of the puzzle may explain how IL PV-INs are ‘turned off’ to support fear extinction encoding, and could potentially implicate dysfunctional IL SST-IN activity in processes underlying extinction deficits and/or fear relapse. Indeed, ventral hippocampal SST-IN activity is critical for shaping contextual fear extinction as well as its relapse^45^ highlighting a role for these cells in other extinction processes in other brain regions.

The plasticity mechanisms underlying recruitment of IL PNs has been studied by a number of groups, and both intrinsic excitability and synaptic plasticity changes have been observed after fear extinction. Specifically, IL PNs exhibit decreased intrinsic excitability after conditioning that is reversed upon extinction; at the same time, IL PNs also showed increased burst firing after extinction training^46,47^. These observations appear to be projection-specific as IL PNs projecting to nucleus accumbens exhibit the reduced excitability phenotype following conditioning that reverses after extinction, whereas IL PNs projecting to BLA exhibited enhanced intrinsic excitability after extinction^24^. Aside from intrinsic excitability changes, a number of lines of evidence exist suggesting that IL PNs undergo synaptic plasticity following fear extinction. Interestingly, most of these studies tie this plasticity to postsynaptic mechanisms and focus on excitatory glutamatergic plasticity^47–52^. In our current study, we find that fear conditioning and extinction induce distinct patterns of presynaptic plasticity in the IL. In particular, conditioning drives enhanced glutamatergic drive from BLA onto IL PV-INs, whereas extinction reverses these effects and drives an additional enhancement of GABAergic drive from IL SST-INs onto IL PV-INs. We also observe enhanced glutamatergic drive from BLA onto IL SST-INs after fear extinction training. Thus, we reveal a potential disinhibitory circuit mechanism driving the recruitment of IL PNs, in addition to their direct innervation by regions like BLA and ventral hippocampus. Our observations of enhanced presynaptic plasticity are unique compared to other studies. One explanation is that our study focuses on local GABAergic circuits, whereas most other studies employing electrophysiological approaches focus on PNs. Moreover, many studies examining synaptic plasticity in IL PNs focus on excitatory mechanisms, likely overlooking some local microcircuit influences that we observed.

Foundational experiments found that manipulations of IL during extinction training impacted extinction retrieval the next day, but had no within-session effects on behavior^10–15^. In our current study, we found that whether we manipulated IL SST-INs or IL PV-INs during extinction training, we observed behavioral effects both within the extinction training sessions as well as during extinction memory retrieval the next day, when cells were not manipulated. These differences could potentially reflect functional roles for these cells that may be distinct from that of PNs. They could be also be attributed to the method by which the cells were manipulated. As our behavioral readouts were freezing bouts, we considered the possibility that our results could potentially be impacted by changes in ambulation that don’t necessarily reflect fear memory states. Indeed, a recent threat avoidance study revealed that IL PV-IN activity is highly correlated with ambulation in this task where mice must actively suppress freezing^53^. To test specificity of IL SST-IN and PV-IN activity to fear/extinction-related behaviors, we performed pre-conditioning/ naïve as well as open field optogenetic tests and did not observe effects of manipulation on locomotion. We additionally quantified calcium traces during freezing onset/offset epochs outside CS presentation and did not detect significant changes in activity of either cell type. These differences in IL PV-IN engagement during suppression of freezing could reflect differences in information processing in each task (spatial information, action-outcome, etc).

After recruitment, IL PNs projecting to BLA selectively exhibit enhanced intrinsic excitability, and chemogenetic inhibition of this projection pathway suppresses extinction memory formation^21,23,24^. Within the BLA, distinct PNs are engaged after fear conditioning compared to fear extinction, and these populations can be distinguished by their projections to PL and IL, respectively^39^. In agreement, we found that BLA engages IL SST-INs in response to fear extinction, and that these BLA->IL projections are required for fear extinction encoding and the associated learning-related plasticity. However, we also found that BLA projections onto IL PV-INs are strengthened after fear conditioning, potentially driving feedforward inhibition onto IL PNs. Thus, in addition to anatomical definitions^39^, BLA projections also seem to exhibit cell type-specific engagement within IL. Such cell type-specific wiring could represent an important circuit mechanism not only for engaging distinct brain regions, but also for engaging specific microcircuits in each region to drive the promotion or suppression of fear. Whether the same BLA projections that target PL PNs and SST-INs^26,27^ also target IL PV-INs to coordinate pro-fear activity profiles across mPFC subregions remains an important area of future investigation.

Overall, we resolved the IL microcircuit connectivity and plasticity mechanisms that support encoding of auditory fear extinction memory. Specifically, we revealed that the engagement of specific GABAergic cell types within the IL microcircuit is a critical mechanism for switching between high and low fear states necessary for fear extinction encoding and expression. Thus, our results highlight how local circuit computations are critical for shaping overall neural activity and behavioral outputs in emotional memory processes.

## Methods

### Animals

The use of animals in all experimental procedures was approved by the Institutional Animal Care and Use Committee at the Heersink School of Medicine at the University of Alabama at Birmingham. Both male and female mice at the age of postnatal day 50 (P50) were used for stereotaxic surgery, and behavioral experiments were then performed at P60-90. Animals were housed in groups of 2-6 with a 12 h light-dark cycle and *ad libitum* access to food and water. An equal number of males and females was used for all experimental groups in all figures. Mice were purchased from The Jackson Laboratory and bred and housed at UAB. Wild-type C57BL/6J mice (Stock No. 000664) and several transgenic lines that were maintained homozygous on a C57BL/6J background, including SST-IRES-Cre (SST-Cre; Stock No. 028864), PV-IRES-Cre (PV-Cre; Stock No. 017320), SST-IRES-FlpO (SST-FlpO; Stock No. 028579), Ai9 (Stock No. 007909), Ai65F (Stock No. 032864), and Ai3 (Stock No. 007903) were used. Experimental mice were generated by crossing these lines as needed to achieve cell-type-specific expression of reporters and actuators. Littermates were equally but randomly assigned to experimental groups/ conditions across all experiments.

### Vectors

Vectors used in our study were purchased from Addgene except where noted. These include AAV1-DIO-eYFP (Cat. No. 27056), AAV8-hSyn-mCherry (Cat. No. 114472), AAV1-syn-FLEX-jGCaMP8f (Cat. No. 162379), AAV1-DIO-hChR2(H134R)-eYFP (Cat. No. 20298), AAV1-FLEXFRT-hChR2(H134R)-mCherry (Cat. No. 75470), AAV1-fDIO-hChR2(H134R)-eYFP (Cat. No. 55639), AAV1-FLEX-Arch-GFP (Cat. No. 22222), AAV1-fDIO-eYFP (Cat. No.55641), AAV1-Ef1a-DIO-ChETA-EYFP (Cat. No. 26968), AAV1-Coff/Fon-Arch3.3-p2a-eYFP (Cat. No. 137150), AAV1-CaMKIIa-hChR2(H134R)-eYFP (Cat. No. 26969), AAV1-CaMKIIa-eYFP (Cat. No. 50469), and AAV1-CaMKIIa-Arch-GFP (Cat. No. 99039). The ESARE-ERCreER plasmid was a gift from Dr. H. Bito (U. Tokyo) and was purified and custom packaged in AAV8 serotype at the Boston Children’s Research Vector Core.

### Stereotaxic surgery

Following induction of general surgical anesthesia, mice (P50) were mounted in stereotaxic frames. For all vector infusions targeting the infralimbic cortex (IL), the following coordinates were used (relative to Bregma): AP: +1.70 mm, ML: ±0.90 mm (at a 15° angle), and DV: -2.3 mm. For activity-dependent neural tagging experiments (Fig. 2), C57Bl/6J received bilateral IL infusions of a viral cocktail containing AAV8-ESARE-ERCreER-PEST, AAV1-DIO-eYFP, and AAV8-hSyn-mCherry, mixed in a 2:7:1 ratio just prior to infusion. Infusion volumes were 400 nl per hemisphere. Vectors incubated for 3 weeks prior to behavioral training. For *in vivo* fiber photometry (Fig. 3), separate cohorts of SST-Cre and PV-Cre mice received unilateral counterbalanced IL infusions of AAV1-syn-FLEX-jGCaMP8f (300 nl infusion volume). An imaging fiber (400 µm diameter; Doric Lenses) was subsequently implanted immediately above the infusion site at AP: +1.70 mm, ML: ±0.90 mm (at a 15° angle), and DV: -2.2 mm. Vectors incubated for 3 weeks prior to behavioral training. For *in vivo* optogenetics (Fig. 4), SST-Flp and PV-Cre mice received bilateral IL infusions of opsins or control vectors. SST-Flp mice received infusions of AAV1-fDIO-hChR2(H134R)-eYFP, AAV1-Coff/Fon-Arch3.3-p2a-eYFP, or AAV1-fDIO-eYFP. PV-Cre mice received infusions of AAV1-Ef1a-DIO-ChETA-EYFP, AAV1-FLEX-Arch-GFP, or AAV1-DIO-eYFP. Vector infusion volume for all constructs and mouse lines was 300 nl per hemisphere. Following infusions, optic fibers (200 µm diameter; Doric Lenses) were implanted bilaterally immediately above IL AP: +1.70 mm, ML: ±0.90 mm (at a 15° angle), and DV: -2.2 mm. Vectors incubated for 2 weeks prior to behavioral training. For microcircuit electrophysiology recordings (Fig. 6), distinct viral strategies were used in triple-transgenic mice. For experiments where synaptic responses were recorded from IL PV-INs, SST-FlpO/ PV-Cre/ Ai65F mice received bilateral IL infusions of AAV1-DIO-ChR2-eYFP (300 nl per hemisphere). For experiments where synaptic responses were recorded from IL SST-INs, SST-FlpO/ PV-Cre/ Ai3 mice received bilateral IL infusions of AAV1-CAG-FLEXFRT-ChR2(H134R)-mCherry. Vectors incubated for 2 weeks prior to behavioral training and subsequent recording. For BLA→IL plasticity experiments (Fig. 7), SST-Flp/ Ai65F and PV-Cre/ Ai9 mice received bilateral infusions of AAV1-CaMKIIa-hChR2-eYFP bilaterally in BLA (AP: -1.4 mm, ML: ±3.5 mm, and DV: -5.0 mm; 200 nl per hemisphere). Vectors incubated for 2 weeks prior to behavioral training and subsequent recording. For *in vivo* optogenetic silencing of BLA→IL experiments (Fig. 8), SST-Flp/ Ai65F and PV-Cre/ Ai9 mice received bilateral infusions of AAV1-CaMKIIa-Arch-GFP or AAV1-CaMKIIa-eYFP bilaterally in BLA (AP: - 1.4 mm, ML: ±3.5 mm, and DV: -5.0 mm; 200 nl per hemisphere). Following infusions, optic fibers (200 µm diameter; Doric Lenses) were implanted bilaterally immediately above IL AP: +1.70 mm, ML: ±0.90 mm (at a 15° angle), and DV: -2.2 mm. Vectors incubated for 2 weeks prior to behavioral training and subsequent recording. For all vector infusions, infusions occurred at a rate of 100 nl/ min. After the conclusion of infusions, needles were left in place for an additional 10 min before being slowly retracted out of the brain.

### Behavioral training paradigms

All behavioral procedures were conducted in sound-attenuating chambers (Med Associates), as previously described^19,26,27^. Across experiments, auditory fear conditioning consisted of six pairings of a conditioned stimulus (CS; 2 kHz, 80 dB, 20 s pure tone) with a co-terminating unconditioned stimulus (US; 0.7 mA, 2 s scrambled foot shock), delivered with an 80 s inter-trial interval. The conditioning context (Context A) consisted of square metal walls and a metal grid floor with 70% ethanol as an odor cue. Extinction training consisted of 1-3 consecutive training days where 20 CS (2 kHz, 80 dB, 20 s pure tone) were presented in a neutral context (Context B) consisting of white, round walls, white solid floor, and 70% isopropanol as an odor cue. Tones exposure tests consisted of 6 CS presentations (2 kHz, 80 dB, 20 s pure tone) in Context A in the absence of US presentation. Retrieval and tones re-exposure tests were performed 24 hours after the conclusion of training and consisted of 4 CS presentations (2 kHz, 80 dB, 20 s pure tone) in Context B. Mice assigned to naïve groups were only experienced to their home cage. Specific training and testing protocols varied according to the experimental goal as follows. Freezing, or the cessation of all movement aside from respiratory activity, was used as the primary behavioral readout for fear or extinction memory.

For cFos immunohistochemistry experiments (Fig. 1), mice were assigned to one of three groups: Tones, Conditioning, or Extinction. The Tones group received six CS presentations in the absence of the US in Context A, followed by a CS re-exposure test 24 h later in Context B. The Fear group underwent fear conditioning in Context A followed by a CS-evoked fear retrieval test 24 h later in Context B. The Extinction group underwent fear conditioning in Context A, followed by two consecutive days of extinction training in Context B, followed by a CS-evoked extinction retrieval test in Context B at 24 h after the last extinction session. Mice were subjected to transcardial perfusion at 90 minutes after the final behavioral test for cFos immunohistochemical analysis (described below).

For activity-dependent neural tagging experiments (Fig. 2), mice were randomly split into one of three groups: fear retrieval, early extinction, or late extinction groups. The fear retrieval group was subjected to fear conditioning in Context A followed by CS-evoked fear retrieval in Context B 24 hours later. The early extinction group was subjected to fear conditioning in context A followed by one day of extinction training in Context B. The late extinction group was subjected to fear conditioning in context A followed by three consecutive days of extinction training in Context B. Mice received a single intraperitoneal injection 4-hydroxytamoxifen (4-OHT; 10 mg/kg) immediately following the end of the last behavior session for each group, as we have done previously^19,27^. Histological analysis was performed at two weeks after tagging to allow for sufficient reporter protein expression.

For *in vivo* fiber photometry calcium imaging experiments (Fig. 3), mice were habituated to handling and imaging cord tethering for 3 days. Mice underwent a within-subject longitudinal protocol, including a pre-conditioning tone exposure test in context B, fear conditioning in context A, a CS-evoked fear retrieval test in context B, two consecutive days of extinction training in context B, and a CS-evoked extinction retrieval test in context B. Calcium signals were continuously recorded throughout all behavioral testing.

For *in vivo* optogenetics experiments (Fig. 4), mice were habituated to handling and patch cord tethering for 3 days, and were tethered in all behavioral tests regardless of whether they received stimulation or not. Mice were first subjected to a tone exposure test consisting of a baseline period with and without counterbalanced 20 s laser manipulation epochs in addition to four CS presentation epochs counterbalanced with and without laser manipulation in context B (stimulation parameters described in *in vivo* optogenetics section). The next day, mice were subjected to fear conditioning without optogenetic manipulation in context A. At 24 hours later, mice were subjected to a two-day extinction training paradigm where optogenetic manipulations were delivered specifically during each CS presentation in context B. A CS-evoked extinction retrieval test was performed 24 h after the final extinction session in context B in the absence of optogenetic manipulation.

For *ex vivo* acute whole-cell brain slice electrophysiology experiments (Fig. 5-8), mice were assigned to one of three groups: naïve, fear, and extinction. Mice in the naïve group remained in their home cage with no behavioral manipulation. Those in the fear group underwent auditory fear conditioning in context A. Mice in the extinction group were subjected to fear conditioning in context A followed by two consecutive days of extinction training in context B. For the experiment described in Fig. 8, which involved optogenetic silencing of BLA projections, animals were subjected to fear conditioning and two subsequent days of extinction training during which BLA projections to IL were optogenetically silenced (see optogenetics methods for laser delivery parameters). For all electrophysiology cohorts, acute brain slices were prepared 24 h after the final behavioral manipulation.

### Activity-dependent neural tagging

Neural tagging was performed as previously described^19,27^. Immediately after CS-evoked fear memory retrieval, day 1 of extinction training, or day 3 of extinction training, mice received a single intraperitoneal injection of 4-hydroxytamoxifen (4-OHT; 10 mg/kg). 4-OHT was formulated in an aqueous medium and delivered as described previously^19,27,54^. Mice were returned to their home cage and left undisturbed for 24 hours. Histology was examined at 2 weeks post-4-OHT tagging.

### Fiber photometry recording and analysis

Fiber photometry recordings were performed using a complete system from Tucker-Davis Technologies, following our previously established protocols^19,26^. At 3 weeks after vector infusion and imaging fiber implantation (0.4 mm core; 0.48 numerical aperture), mice were habituated to handling and imaging cord (0.4 mm core; 0.48 numerical aperture) tethering for 3 consecutive days. Calcium signals were monitored during handling and habituation to ensure reliable and stable signals prior to behavioral testing. LED light at 465 nm was used to monitor calcium-associated fluorescence fluctuations while 405 nm light was simultaneously used as an isosbestic control for motion artifacts. Light was delivered into the brain at a final transmittance of 40-80 µW and kept constant across days. Calcium signals were sampled at 6 kHz and continuously recorded throughout each behavioral test. Transistor-transistor logic (TTL) signals were generated through FreezeFrame (MedAssociates) software to produce timestamps for session boundaries and CS presentation epochs. AnyMaze (Stoelting) was used to automatically detect and generate TTL signals for freezing onset and offset epochs during baseline, CS, and inter-CS intervals. Prior to the start of each behavioral session, calcium signals were recorded for 2 min to ensure stable recordings.

Data were processed and analyzed using custom-written code as described previously^19,26^. Briefly, CS-associated activity changes were assessed by normalizing the calcium signal during the 20 sec CS presentation (or 18 sec for conditioning sessions) to a 20 sec baseline period (or 18 sec for conditioning sessions) occurring immediately prior to CS onset. The percentage change in peak fluorescence was calculated as %ΔF/F = (FpeakCS - FpeakBL)/FpeakCS. Comparisons in fluorescence changes across days was normalized to CS-evoked signals recorded during the pre-conditioning CS exposure test. Reported values represent within animal changes.

### In vivo optogenetics

Two weeks following vector infusion and optic fiber implantation, mice were habituated to handling and patch cord (200 µm diameter) tethering for 3 consecutive days. For ChR2-expressing IL SST-INs, light (465 nm) was delivered via 10 ms pulses at 20 Hz for 20 sec epochs. For Arch-expressing IL SST-INs, solid light (532 nm) was delivered for 20 sec epochs and ramped down for a period of 5 seconds at the conclusion of CS to prevent rebound action potentials. For ChETA-expressing IL PV-INs, light (465 nm) was delivered via 2 ms pulses at 50 Hz for 20 sec epochs. For Arch-expressing IL PV-INs, solid light (532 nm) was delivered for 20 sec epochs and ramped down for a period of 5 seconds at the conclusion of CS to prevent rebound action potentials. eYFP control groups for SST-ChR2, SST-Arch, PV-ChETA, and PV-Arch all received the same light pattern deliveries as their opsin-containing counterparts. Light was delivered in the brain at a final transmittance of 6-9 mW for all opsin groups. To evaluate potential light-induced behavioral changes occurring prior to behavioral training, mice underwent a pre-conditioning baseline testing session consisting of light-on and light-off epochs in the absence and presence of CS presentation in a counterbalanced manner (e.g. Baseline: light-on, light-off; CS: CS + light, CS alone, etc) and the order of light presentation was alternated. Each epoch was repeated twice and behavioral readouts were reported as an average of the two epochs. Mice then underwent paired auditory fear conditioning while tethered to patch cords, but in the absence of optogenetic manipulation. Subsequently, mice underwent extinction training in context B over two consecutive days, during which optogenetic activation or silencing was delivered concurrently with each CS presentation. A CS-evoked extinction memory retrieval test was conducted 24 h later in context B in the absence of optogenetic manipulation. Behavioral videos were scored by experimenters blind to groups, conditions, and stimulation epoch order. Following experiments, brains were processed for histological verification of fiber placement and opsin expression, and mice with off-target fiber placements or vector mistargeting were excluded from analysis.

### Open field test

Mice tethered to patch cords were placed in a 40 cm x 40 cm x 40 cm (l x w x h) white opaque open field arena. The 10-minute session was divided into alternating 1-minute intervals of light-on and light-off periods, with the starting condition counterbalanced among subjects. Stimulation parameters for each opsin and control group were identical to those used for stimulation during behavioral training. Mice were tracked from an overhead perspective and analyzed using EthoVision XT tracking software (Noldus). Total distance traveled and mean velocity were quantified for each epoch for each mouse. Each of the light-on (total of 5) and light-off (total of 5) epochs were averaged and reported as single values for each mouse.

### Whole-cell brain slice electrophysiology

At 24 hours after the last behavioral training session, mice were deeply anesthetized, decapitated, and brains removed. Acute brain slices (350 µm) containing the infralimbic cortex were prepared using a Leica VT1200S vibratome in a chilled, carbogen-bubbled (95% O_2_ and 5% CO_2_), sucrose cutting solution containing (in mM) 210 sucrose, 11 glucose, 26.2 NaHCO_3_, 2.5 KCl, 1 NaH_2_PO_4_, 4 MgCl_2_, 0.5 CaCl_2_, and 0.5 ascorbate. Slices recovered for 45 minutes at 34°C in carbogen-bubbled artificial cerebrospinal fluid (ACSF) containing (in mM) 119 NaCl, 26.2 NaHCO_3_, 11 glucose, 1 NaH_2_PO_4_, 2 MgCl_2_, 2.5 KCl, 2 CaCl_2_. Slices were kept at room temperature for the duration of the recordings. Whole-cell patch-clamp recordings were made using electrodes (3-4 MΩ) filled with a cesium-methanesulfonate internal solution containing (in mM) 120 Cs-methanesulfonate, 10 HEPES, 10 Na-phosphocreatine, 1 QX-314, 8 NaCl, 0.5 EGTA, 4 Mg-ATP, 0.4 Na-ATP (pH 7.25; 290–300 mOsmol) for voltage-clamp experiments. Cells were visualized on an upright microscope with differential interference contrast optics. SST- and PV-expressing interneurons were identified in slices from reporter mice (SST-FlpO/ Ai65F and PV-Cre/ Ai9, respectively) by visualizing their native tdTomato fluorescence under illumination. In contrast, principal neurons (PNs) were identified by their large pyramidal soma and confirmed by their distinct electrophysiological signature (low membrane resistance <70 MΩ; high capacitance >100 pF). Miniature excitatory and inhibitory postsynaptic currents (mEPSCs and mIPSCs, respectively) were recorded (Fig. 3) in ACSF containing 1 µM tetrodotoxin (TTX; HelloBio). Currents were isolated by clamping the cell at -60 mV (for mEPSCs) or 0 mV (for mIPSCs), and each recording lasted a minimum of 5 minutes. For experiments involving optogenetically-evoked synaptic currents (Fig. 6 and 7), light pulses from an LED coupled to the slice microscope objective lens. Short (1 ms) light pulses were used to stimulate neurotransmitter release from specific ChR2-expressing sources (IL SST-INs, IL PV-INs, or BLA afferents in IL) in ACSF with 1 µM TTX and 100 µM 4-aminopyrimidine (Abcam). To compare apparent synaptic strength across behavioral conditions, the amplitudes of light-evoked currents recorded from interneurons were normalized to the average amplitude of currents recorded from neighboring PNs within the same slice. For optogenetics-assisted paired-pulse experiments, pairs of light pulses at varying inter-stimulus intervals were delivered. Paired pulse ratio was calculated by dividing the amplitude of the second pulse by the amplitude of the first pulse. Data were acquired at 10 kHz and low-pass-filtered at 3 kHz using Multiclamp 700B amplifier. All recordings were analyzed offline by experimenters blind to the experimental conditions using Easy Electrophysiology (v2.8.0) or Clampfit 11.3 (Molecular Devices). Recordings were discarded if access resistance changed by >20% or if the leak current was unstable or exceeded 200 pA during recording.

### Immunohistochemistry

For all histological analyses, at the conclusion of experiments, mice were subjected to transcardial perfusion using ice-cold phosphate-buffered saline (PBS; pH 7.4) followed by 4% paraformaldehyde (PFA) in PBS. Brains were extracted and post-fixed in 4% PFA overnight at 4°C, then sectioned coronally at 50 µm using a Leica VT1000S vibratome. For all experiments, free-floating sections were blocked for one hour at room temperature in PBS containing 5% normal bovine serum albumin and 0.25% Triton X-100 (Sigma). For cFos immunohistochemical analysis (Fig. 1), slices were subsequently incubated overnight at 4°C with a rabbit anti-cFos antibody (1:1000; Millipore, ABE457) in PBS containing 2.5% normal bovine serum albumin and 0.25% Triton X-100. The following day, sections were thoroughly washed in PBS and incubated for two hours at room temperature with a goat anti-rabbit secondary antibody conjugated to Alexa Fluor 488 (1:500; Jackson ImmunoResearch) in PBS containing 2.5% normal bovine serum albumin and 0.25% Triton X-100. To determine the neurochemical identity of neurons recruited by distinct behavioral states (Fig. 2), parallel sets of sections from the same mice were processed in two experiments. First, a subset of sections was incubated overnight at 4°C with a rabbit anti-somatostatin antibody (1:1000; BMA Biomedicals, T-4103) in PBS containing 2.5% normal bovine serum albumin and 0.25% Triton X-100. A second set of sections was incubated with a mouse anti-parvalbumin antibody (1:1000; Sigma-Aldrich, MAB1572). The following day, after thoroughly washing sections with PBS, the respective sections were incubated with their corresponding secondary antibodies (goat anti-rabbit conjugated to Alexa 647 (1:500; Jackson ImmunoResearch) or goat anti-mouse conjugated to Alexa 647 (1:500; Jackson ImmunoResearch)). To amplify the GFP/eYFP signal for all experiments for the purpose of validating viral expression and enhancing reporter visualization (Figs. 2,3,6,7,8), sections were incubated overnight at 4°C with a chicken anti-GFP primary antibody (1:500; Millipore, AB16901) in PBS containing 2.5% normal bovine serum albumin and 0.25% Triton X-100. The following day, sections were thoroughly washed with PBS and incubated for two hours at room temperature with a donkey anti-chicken secondary antibody conjugated to Alexa Fluor 488 (1:500; Jackson Immunoresearch). Following immunohistochemical staining, brain sections were mounted onto slides using ProLong Gold antifade reagent with DAPI. Imaging was performed using a Nikon Eclipse Ni-E microscope, and images were acquired as z-stacks with a 5 µm step size.

### Quantification of immunohistochemical images

Following image acquisition, anatomical demarcations of the infralimbic cortex were determined by creating digital overlays aligned to Paxinos and Franklin’s The Mouse Brain in Stereotaxic Coordinates atlas^55^ in Adobe Photoshop. Cell counting was performed manually using the Cell Counter plugin in ImageJ (NIH) by observers blind to experimental groups and conditions. The specific quantification varied by experiment. For Figure 1, tdTomato-positive reporter cells and cFos-positive cells were first quantified, followed by subsequent quantification of their co-localization determined by independent marker overlap. For Figure 2, eYFP-tagged neurons and SST- or PV-immunoreactive cells were first quantified, followed by subsequent quantification of their co-localization determined by independent marker overlap. For all other experiments, this methodology was used to verify anatomical location and area of viral vector expression, as well as locations of imaging and optic fiber tip terminations. Mice with incorrect viral targeting or spread and/or incorrect fiber placements were excluded from analysis.

### Statistical analysis

Levine’s and Shapiro-Wilk tests were performed prior to parametric statistical testing to establish whether data exhibited homogeneity of variance and normal distribution, respectively. If either or both conditions were not met, data were subjected to equivalent non-parametric statistical analyses. Statistical testing and graph plotting were performed using OriginPro 2025b version 1.2.5.234.

## Acknowledgments

This work was supported by National Institute on Drug Abuse grant no. R01 DA059368 to K.A.C., National Institute of Mental Health grant no. R00 MH122228 to K.A.C., the James A. Pittman Scholar Award to K.A.C., Joe W. and Dorothy Dorsett Brown Foundation grant to K.A.C., T32 MH129274 to H.T.F., T32 GM146611 to Z.R.D., and T32 NS061788 to B.L.F. Part of this work was supported by Boston Children’s Hospital Viral Core, which is supported by NIH5P30EY012196. We thank Dr. Sofia Beas, Dr. Bryan Luikart, and Dr. Jamie Peters as well as members of the Cummings lab for helpful feedback on the manuscript.

## Author contributions

R.C.C. and K.A.C. initiated the project. K.A.C. supervised the research. K.A.C., R.C.C., H.T.F., and N.R.P. designed experiments. R.C.C., K.A.C., H.T.F., N.R.P., Z.R.D., B.L.F., R.L.F., F.A.B., V.A.L., and M.P. performed the research and analyzed the data. K.A.C. and R.C.C. wrote the manuscript.

## Declaration of interests

The authors declare no competing interests.

---

**Supplementary Figure 1.**
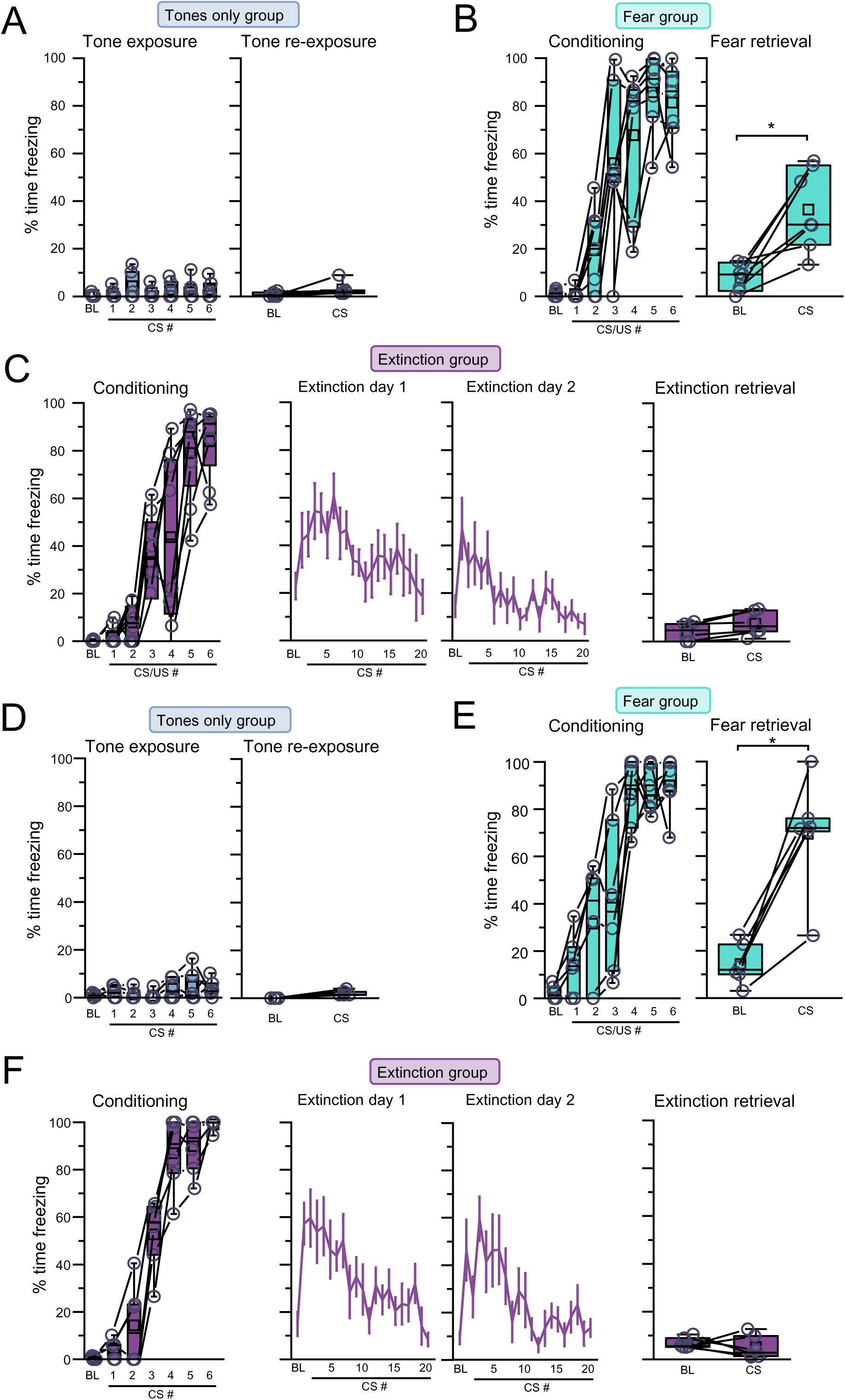
Behavior analysis related to Figure 1. Quantification of freezing for SST-Flp/ Ai65F mice from **(A)** tones (tone re-exposure: W = 3, p = 0.14, Wilcoxon signed ranks test; n = 6 mice), **(B)** fear (fear retrieval: W = 0, p = 0.022, Wilcoxon signed rank test; n = 7 mice), and **(C)** extinction (extinction retrieval: t_5_ = -3.47, p = 0.02, paired t-test; n = 6 mice) groups. Quantification of freezing for PV-Cre/ Ai9 mice from **(D)** tones (tone re-exposure: W = 0, p = 0.06, Wilcoxon signed ranks test; n = 5 mice), **(E)** fear (fear retrieval: W = 14, p = 0.53, Wilcoxon signed ranks test; n = 6 mice), and **(F)** extinction (extinction retrieval: t_5_ = -0.99, p = 0.37, paired t-test; n = 6 mice) groups. Boxplots represent the median (center line), mean (square), quartiles, and 10%–90% range (whiskers/error bars). Open circles represent data points for individual mice.

**Supplementary Figure 2.**
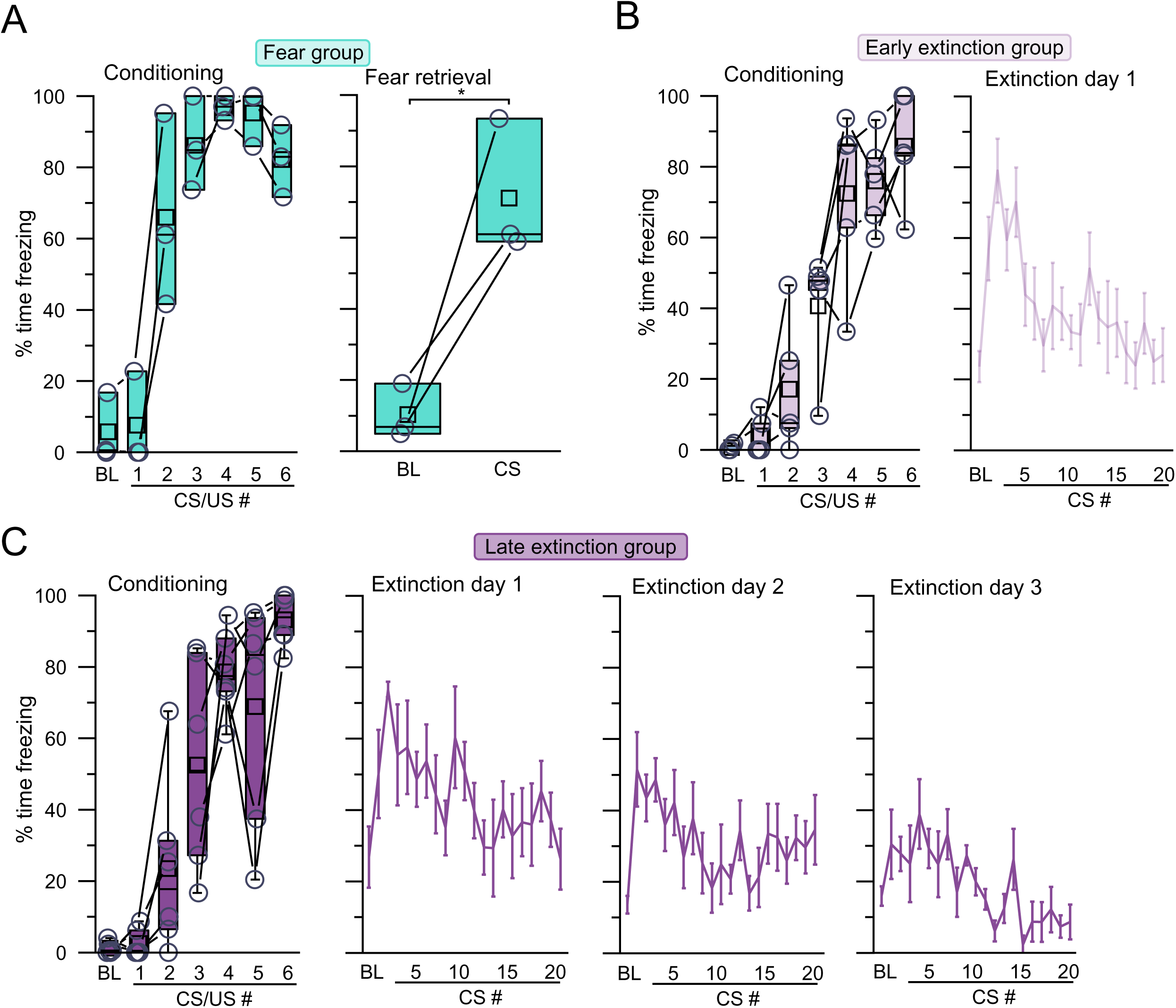
Behavior analysis related to Figure 2. Quantification of freezing for wild-type mice from **(A)** fear (fear retrieval: t_2_ = -4.57, p = 0.04, paired t-test; n = 3 mice), **(B)** early extinction (n = 5 mice), and **(C)** late extinction (n = 6 mice) groups. Boxplots represent the median (center line), mean (square), quartiles, and 10%–90% range (whiskers/error bars). Open circles represent data points for individual mice.

**Supplementary Figure 3.**
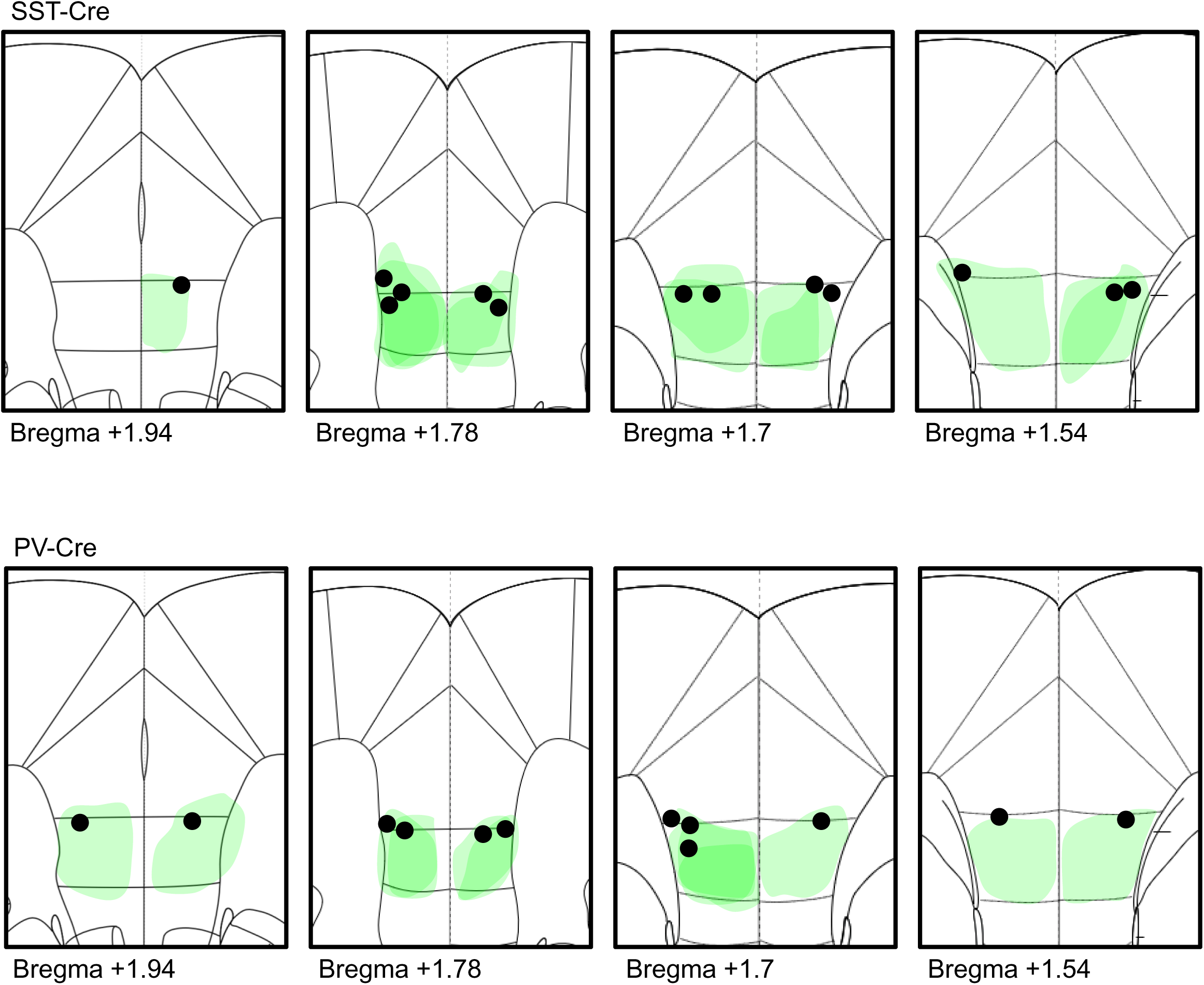
Vector expression and fiber placements related to Figure 3. Cartoon representation of vector spread (green shapes) and optic fiber placements (circles) for **(A)** SST-Cre and **(B)** PV-Cre mice.

**Supplementary Figure 4.**
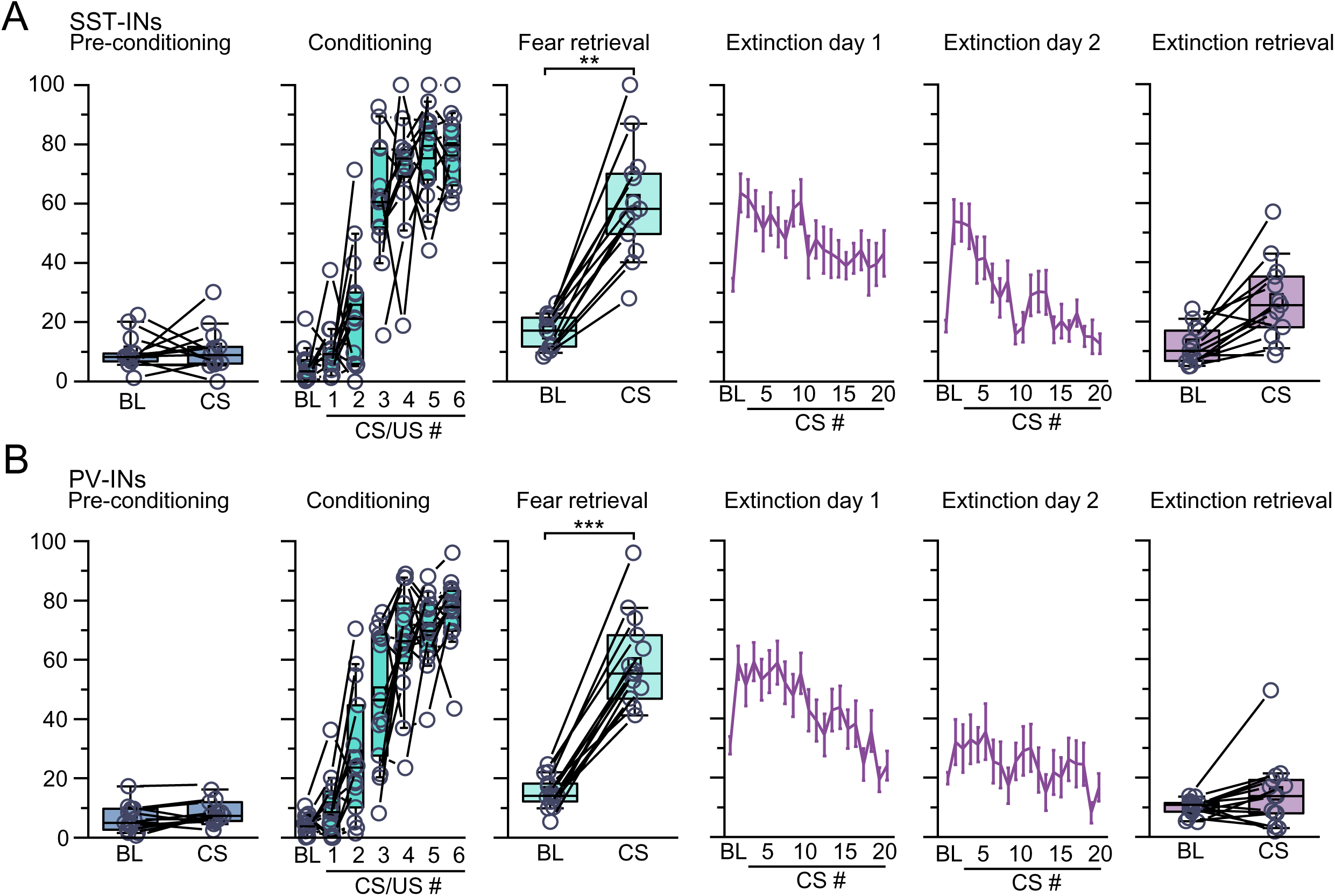
Behavior analysis related to Figure 3. **(A)** Quantification of freezing for SST-Cre mice (n = 13) during pre-conditioning tone exposure (W = 40, p = 0.72, Wilcoxon signed ranks test), conditioning, fear memory retrieval (W = 0, p = 0.002, Wilcoxon signed ranks test), extinction training, and extinction memory retrieval (W = -2, p = 0.051, Wilcoxon signed ranks test). **(B)** Quantification of freezing for PV-Cre mice (n = 15) during pre-conditioning tone exposure (W = 40, p = 0.27, Wilcoxon signed ranks test), conditioning, fear memory retrieval (W = 0, p = 7.3×10^-4^, Wilcoxon signed ranks test), extinction training, and extinction memory retrieval (W = 28, p = 0.13, Wilcoxon signed ranks test). Boxplots represent the median (center line), mean (square), quartiles, and 10%–90% range (whiskers/error bars). Open circles represent data points for individual mice.

**Supplementary Figure 5.**
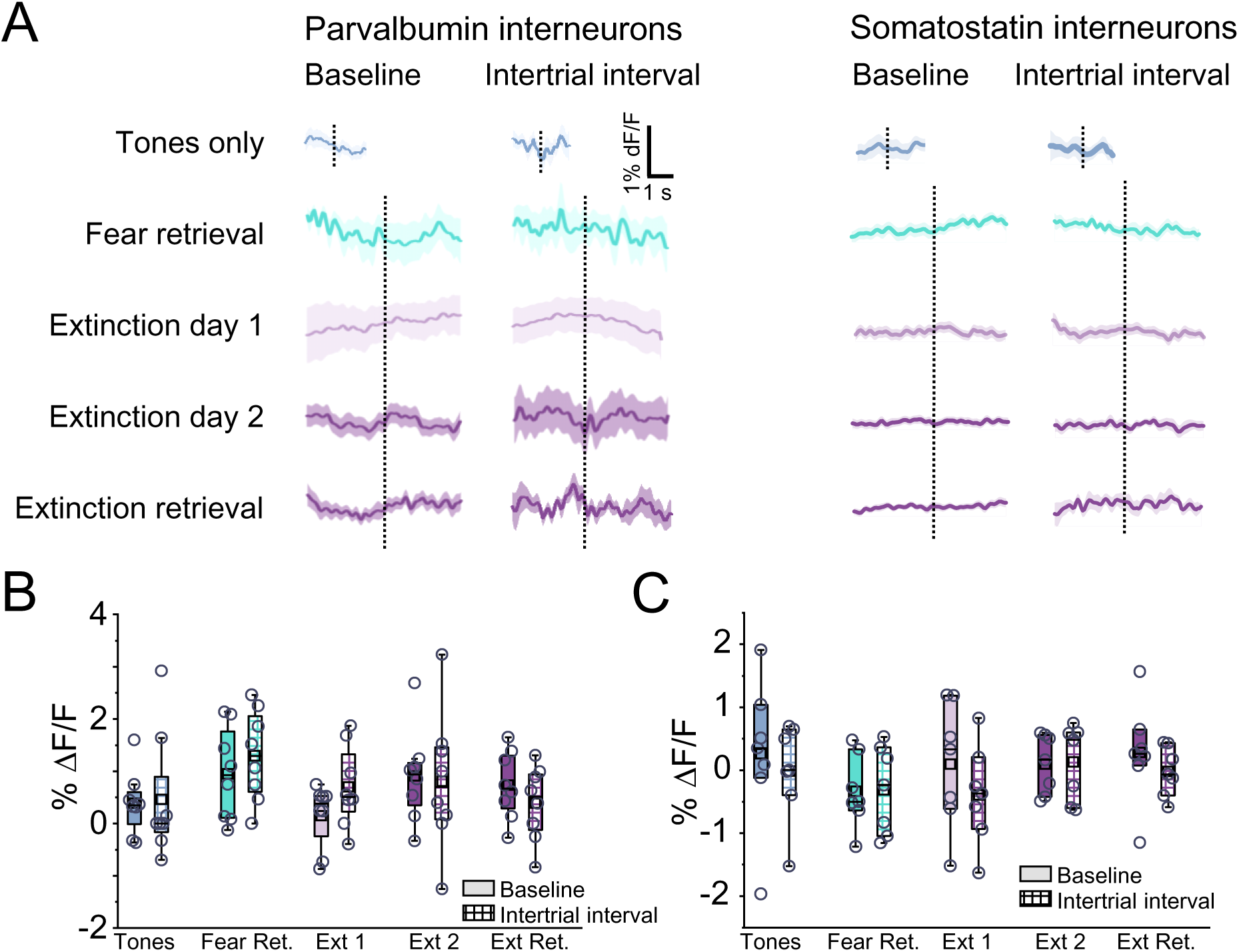
Calcium-associated signals during baseline and intertrial interval related to Figure 3. **(A)** Average calcium-associated signals recorded and aligned for freezing onset during baseline and intertrial interval for infralimbic cortex parvalbumin (left) and somatostatin (right) interneurons across tones only, fear retrieval, extinction days 1 and 2, and extinction retrieval. Criteria for freezing for tones-only were shortened since these epochs were excessively rare. Quantification of freezing-associated calcium signals across behaviors tests during baseline and intertrial interval epochs for **(B)** parvalbumin interneurons (X^2^ = 9.91 [9], p = 0.36, Friedman ANOVA) and **(C)** somatostatin interneurons (F_9,60_ = 0.72, p = 0.69, one-way repeated measures ANOVA). Boxplots represent the median (center line), mean (square), quartiles, and 10%–90% range (whiskers/error bars). Open circles represent data points for individual mice.

**Supplementary Figure 6.**
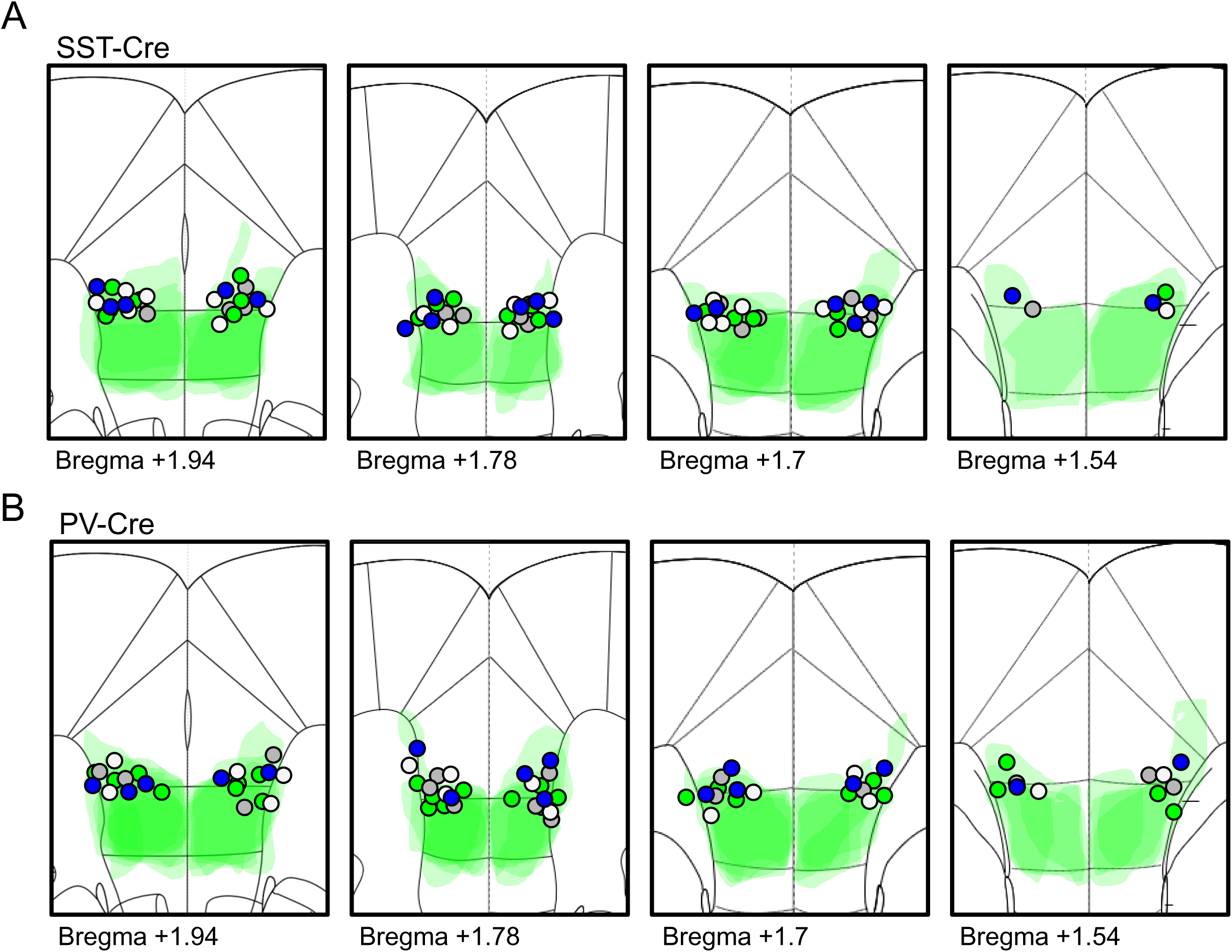
Vector expression and fiber placements related to Figure 4. Cartoon representation of vector spread (green shapes) and optic fiber placements (circles) for **(A)** SST-Flp and **(B)** PV-Cre mice. Gray circles = fiber tip placements for eYFP (paired with Arch mice); green circles = fiber tip placements for Arch; white circles = fiber tip placements for eYFP (paired with ChR2 or ChETA mice); blue circles = fiber tip placements for ChR2 or ChETA.

**Supplementary Figure 7.**
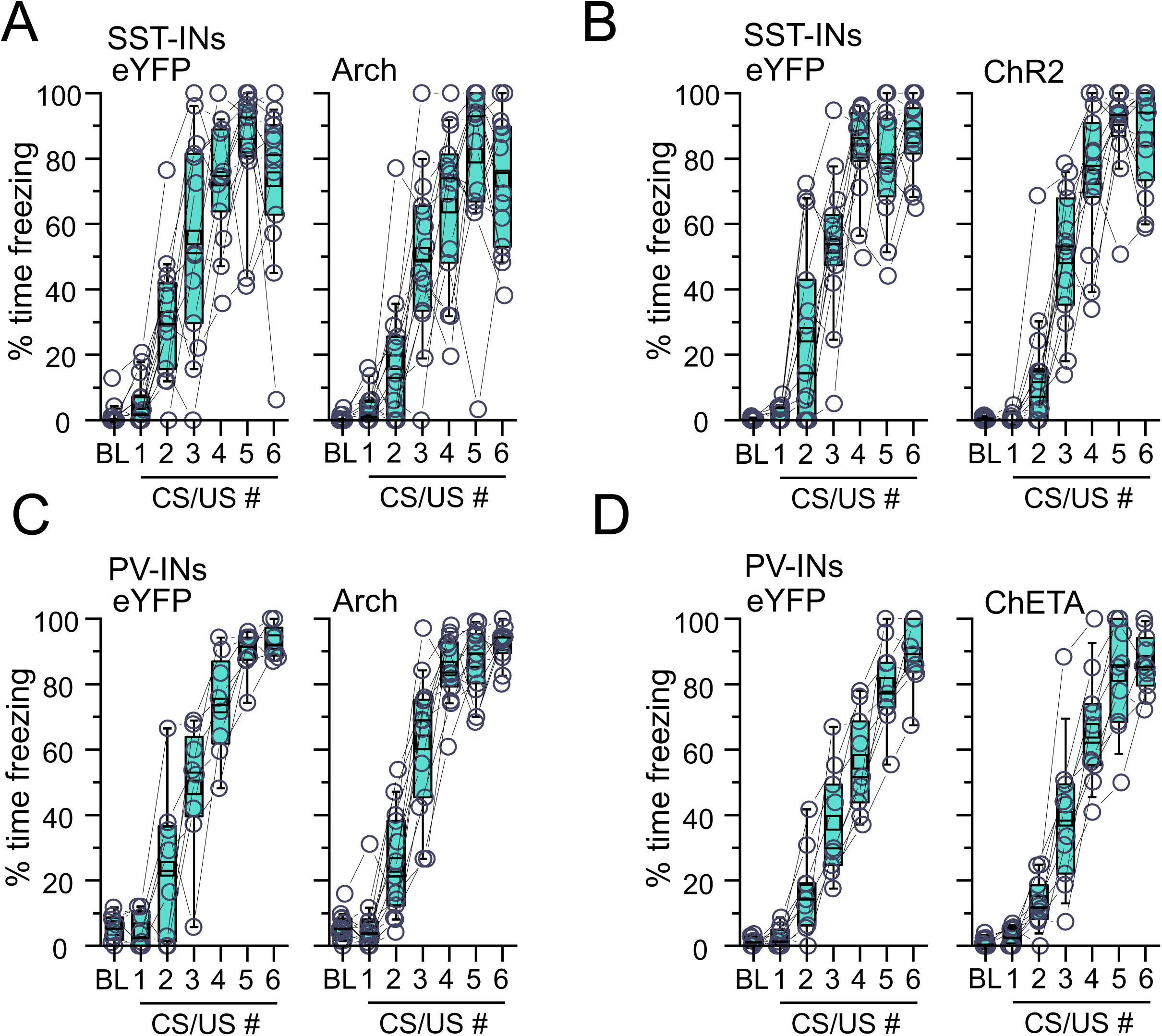
Behavior analysis during conditioning related to Figure 4. Quantification of freezing during fear conditioning for (A) SST-Flp mice expressing eYFP or Arch, (B) SST-Flp mice expressing eYFP or ChR2, (C) PV-Cre mice expressing eYFP or Arch, and (D) PV-Cre mice expressing eYFP or ChETA. Boxplots represent the median (center line), mean (square), quartiles, and 10%–90% range (whiskers/error bars). Open circles represent data points for individual mice.

**Supplementary Figure 8.**
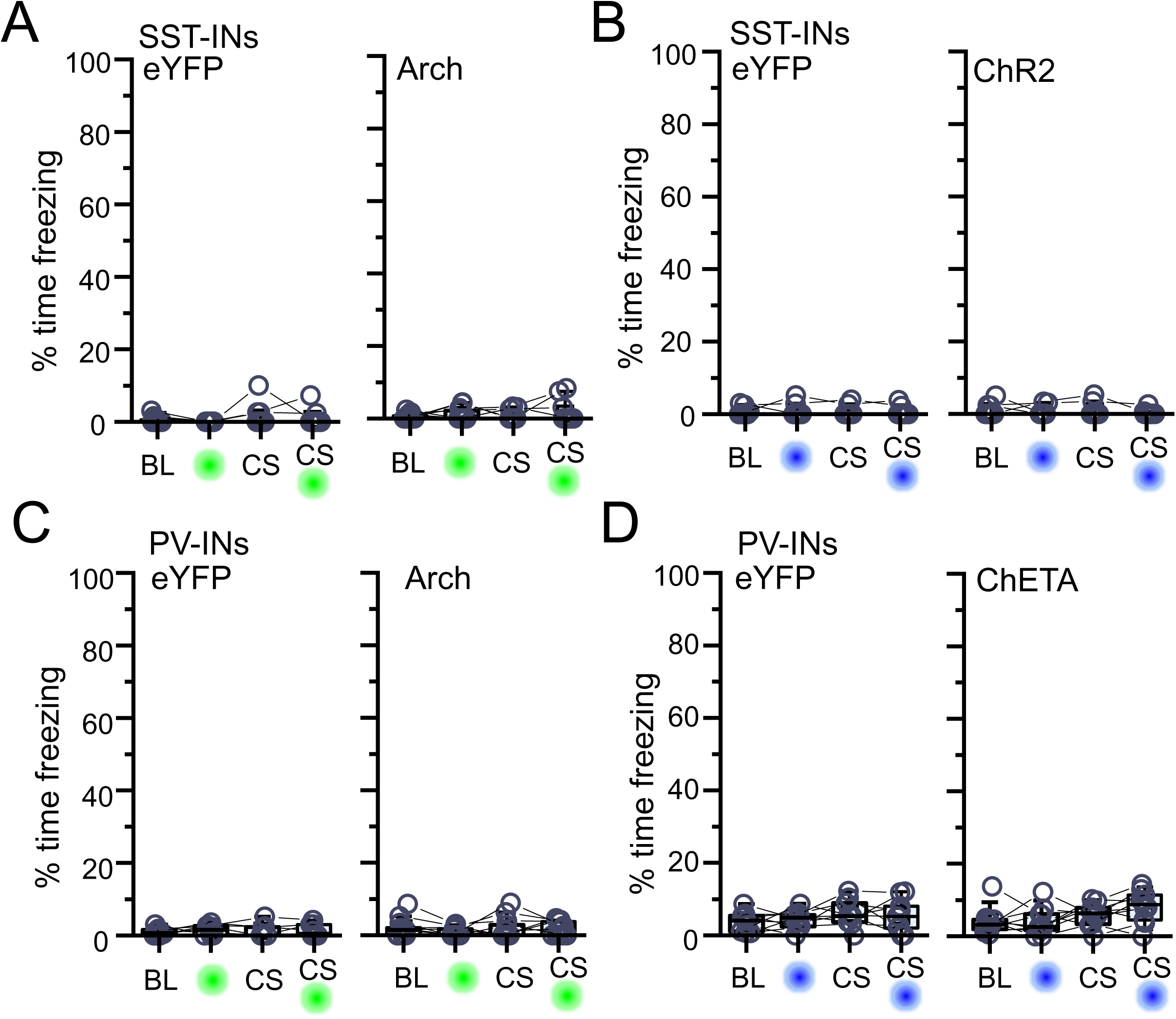
Optogenetic manipulation of infralimbic cortex SST-INs and PV-INs during naïve tone exposure related to Figure 4. Quantification of freezing during baseline, laser stimulation (green or blue circle only), CS presentation, and CS presentation in combination with laser stimulation (CS and green or blue circle) for **(A)** SST-Flp mice expressing eYFP (Χ^2^ = 1.41 [3], p = 0.70, Friedman ANOVA) or Arch (Χ^2^ = 0.13 [3], p = 0.99, Friedman ANOVA), **(B)** SST-Flp mice expressing eYFP (Χ^2^ = 0.28 [3], p = 0.96, Friedman ANOVA) or ChR2 (Χ^2^ = 0.37 [3], p = 0.95, Friedman ANOVA), **(C)** PV-Cre mice expressing eYFP (Χ^2^ = 0.83 [3], p = 0.84, Friedman ANOVA) or Arch (Χ^2^ = 2.38 [3], p = 0.50, Friedman ANOVA), or **(D)** PV-Cre mice expressing eYFP (F_3,24_ = 1.13, p = 0.36, one-way repeated-measures ANOVA) or ChETA (Χ^2^ = 6.87 [3], p = 0.08, Friedman ANOVA). Boxplots represent the median (center line), mean (square), quartiles, and 10%–90% range (whiskers/error bars). Open circles represent data points for individual mice.

**Supplementary Figure 9.**
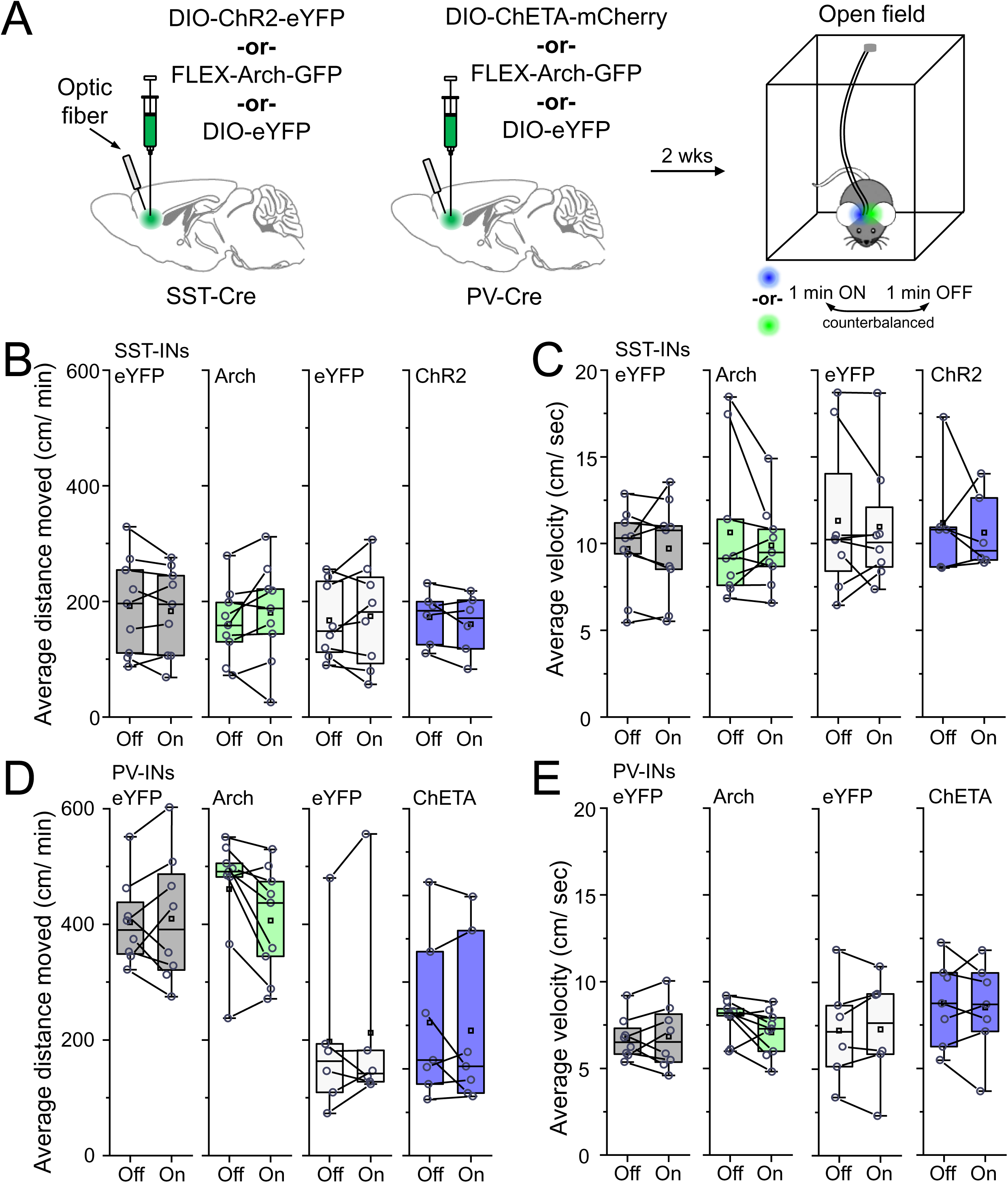
Optogenetic manipulation of infralimbic cortex SST-INs and PV-INs during open field measurements related to Figure 4. **(A)** SST-Flp mice received infusions of FRT-ChR2-eYFP, FRT-Arch-GFP, or FRT-eYFP while PV-Cre mice received infusions DIO-ChETA-eYFP, FLEX-Arch-GFP, or DIO-eYFP. Optical fibers were also implanted and directed at the infralimbic cortex for all mice. After 2 weeks, mice were placed in an open field arena for 10 minutes and were subjected to alternating counterbalanced presentations of optogenetic manipulation. **(B)** Quantification of average distance moved for SST-Flp mice expressing eYFP (paired with Arch mice; t_8_ = 0.96, p = 0.36, paired t-test; n = 9 mice), Arch (t_8_ = -1.36, p = 0.21, paired t-test; n = 9 mice, eYFP (paired with ChR2 mice; t_7_ = -0.52, p = 0.62, paired t-test; n = 8 mice), or ChR2 (t_5_ = 1.41, p = 0.22, paired t-test; n = 6 mice). **(C)** Quantification of average velocity for SST-Flp mice expressing eYFP (paired with Arch mice; t_8_ = -0.06, p = 0.95, paired t-test; n = 9 mice), Arch (W = 26, p = 0.72, Wilcoxon signed ranks test; n = 9 mice), eYFP (paired with ChR2 mice; t_7_ = 0.54, p = 0.61, paired t-test; n = 8 mice), or ChR2 (W = 13, p = 0.67, Wilcoxon signed ranks test; n = 6 mice). **(D)** Quantification of average distance moved for PV-Cre mice expressing eYFP (paired with Arch mice; t_7_ = -0.28, p = 0.79, paired t-test; n = 8 mice), Arch (W = 26, p = 0.06, Wilcoxon signed ranks test; n = 9 mice), eYFP (paired with ChETA mice; W = 7, p = 0.53, Wilcoxon signed ranks test; n = 6 mice), or ChETA (W = 16, p = 0.80, Wilcoxon signed ranks test; n = 7 mice). **(E)** Quantification of average velocity for PV-Cre mice expressing eYFP (paired with Arch mice; t_7_ = -0.28, p = 0.79, paired t-test; n = 8 mice), Arch (W = 39, p = 0.06, Wilcoxon signed ranks test; n = 9 mice), eYFP (paired with ChETA mice; t_5_ = -0.17, p = 0.87, paired t-test; n = 6 mice), or ChETA (t_6_ = 0.49, p = 0.64, paired t-test; n = 7 mice). Boxplots represent the median (center line), mean (square), quartiles, and 10%–90% range (whiskers/error bars). Open circles represent data points for individual mice.

**Supplementary Figure 10.**
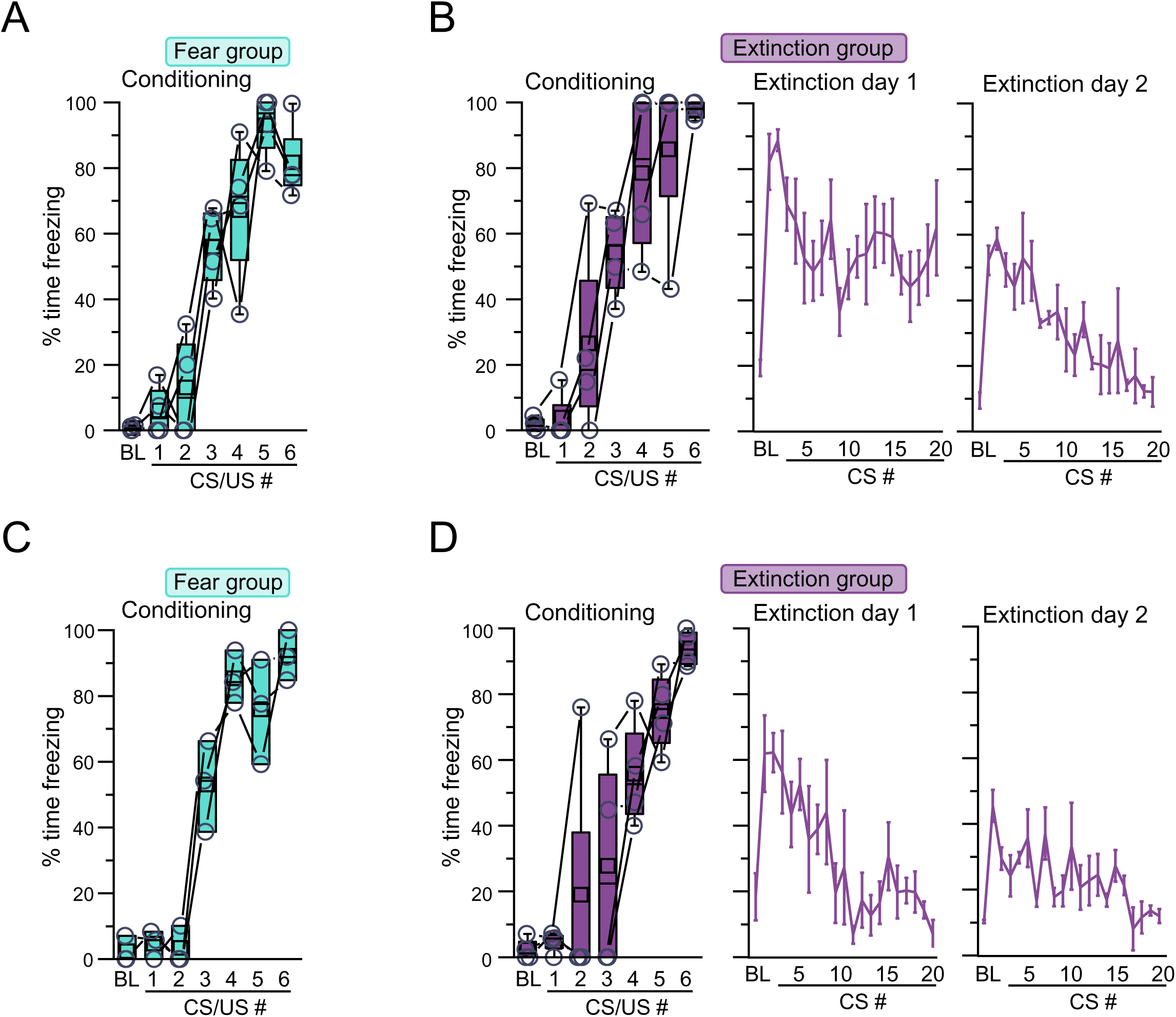
Behavior analysis related to Figure 5. Quantification of freezing for SST-Flp/ Ai65F mice from **(A)** fear (n = 4 mice) and **(B)** extinction (n = 6 mice) groups. Quantification of freezing for PV-Cre/ Ai9 mice from **(C)** fear (n = 3 mice) and **(D)** extinction (n = 4 mice) groups. Boxplots represent the median (center line), mean (square), quartiles, and 10%–90% range (whiskers/error bars). Open circles represent data points for individual mice.

**Supplementary Figure 11.**
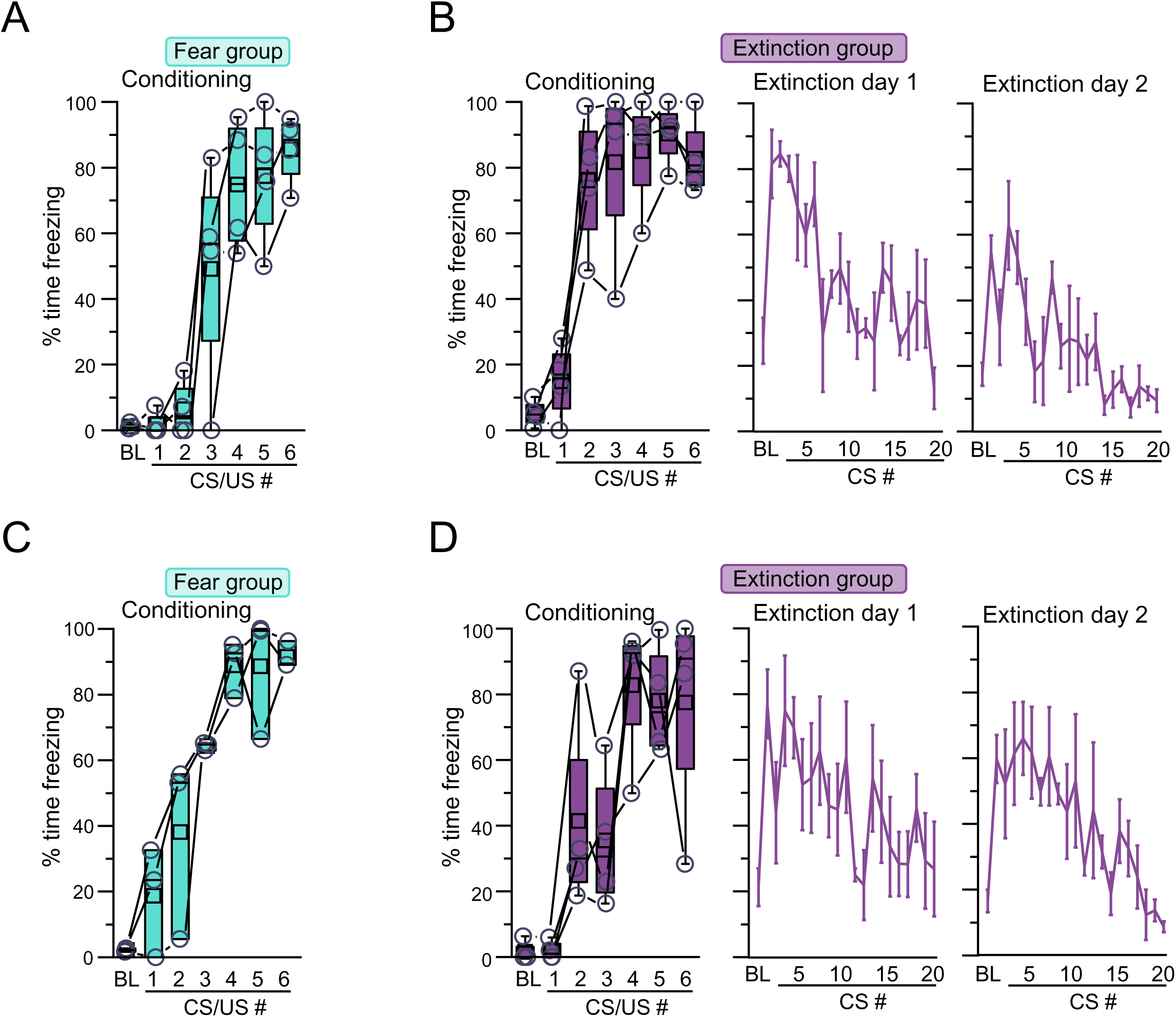
Behavior analysis related to Figure 6. Quantification of freezing for SST-Flp/ PV-Cre/ Ai3 mice infused with FRT-ChR2-mCherry from **(A)** fear (n = 4 mice) and **(B)** extinction (n = 4 mice) groups. Quantification of freezing for SST-Flp/ PV-Cre/ Ai65F mice infused with DIO-ChR2-eYFP from **(C)** fear (n = 3 mice) and **(D)** extinction (n = 4 mice) groups. Boxplots represent the median (center line), mean (square), quartiles, and 10%–90% range (whiskers/error bars). Open circles represent data points for individual mice.

**Supplementary Figure 12.**
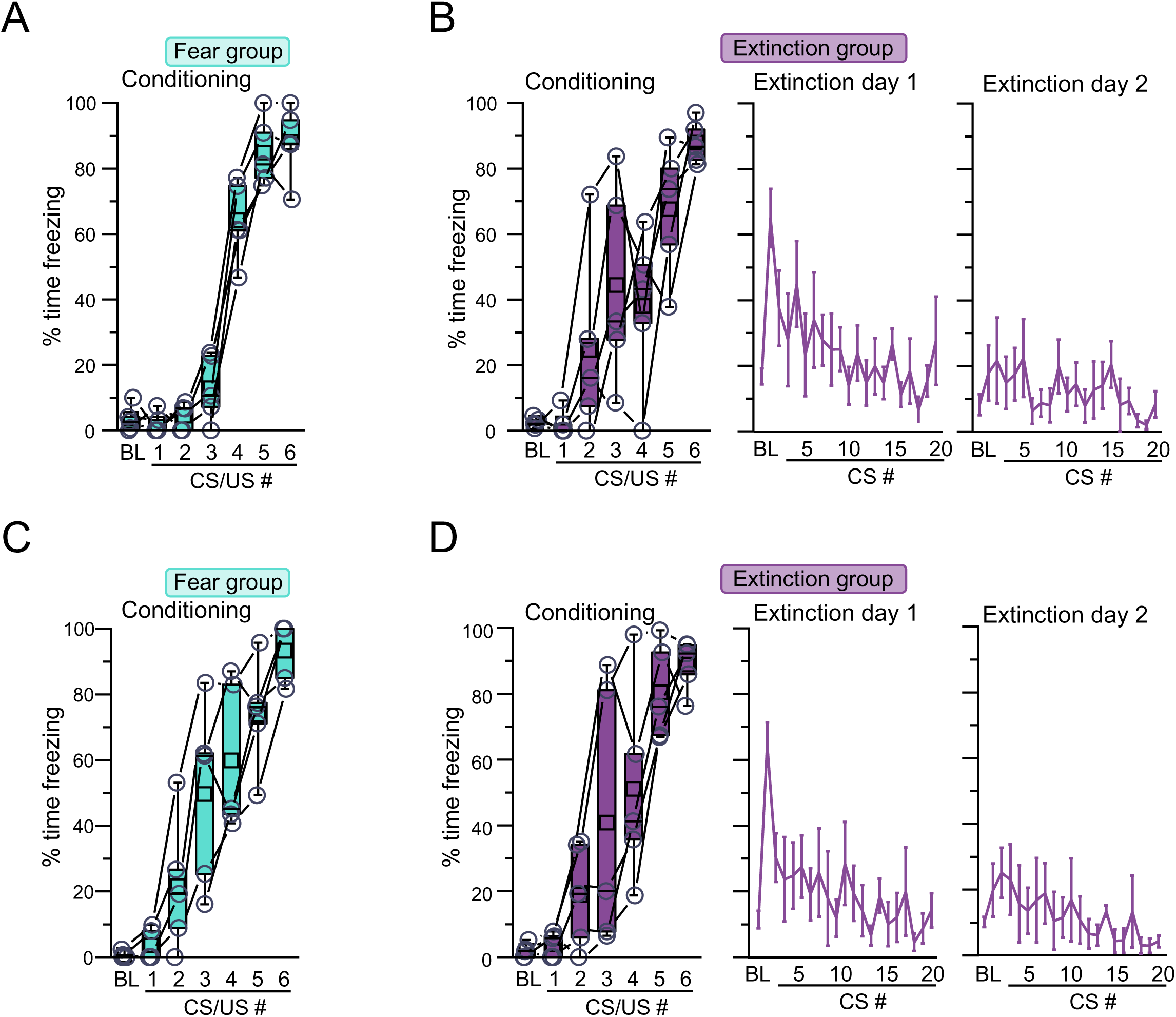
Behavior analysis related to Figure 7. Quantification of freezing for SST-Flp/ Ai65F mice from **(A)** fear (n = 5 mice) and **(B)** extinction (n = 5 mice) groups. Quantification of freezing for PV-Cre/ Ai9 mice from **(C)** fear (n = 5 mice) and **(D)** extinction (n = 5 mice) groups. Boxplots represent the median (center line), mean (square), quartiles, and 10%–90% range (whiskers/error bars). Open circles represent data points for individual mice.

**Supplementary Figure 13.**
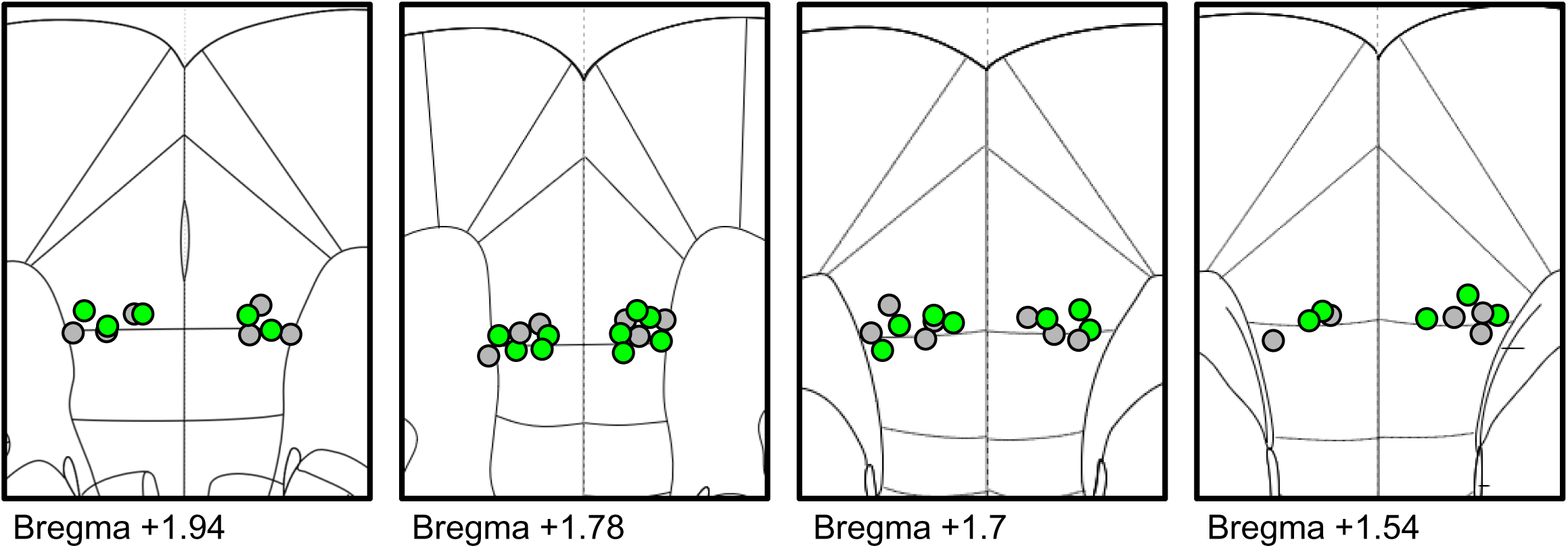
Fiber placements related to Figure 8. Cartoon representation of optic fiber placements (circles) in medial prefrontal cortex. Gray circles = fiber tip placements for eYFP; green circles = fiber tip placements for Arch.

## Notes

### Competing Interest Statement

The authors have declared no competing interest.

